# Minimal eukaryotic DNA polymerase ζ/Rev1 GC-pair mutasome

**DOI:** 10.64898/2026.11.20.768204

**Authors:** Charlie T. McNellis, Morgan Brown, Dmitrii D. Panteleev, Anna S. Zhuk, Igor B. Rogozin, Anna Y. Aksenova, Youri I. Pavlov

**Affiliations:** Eppley Institute for Research in Cancer; Fred and Pamela Buffett Cancer Center, the University of Nebraska Medical Center, Omaha, NE, 68198, USA; College of Saint Mary, Omaha, Nebraska; Institute of Applied Computer Science, ITMO University, 197101, St. Petersburg, Russia; Vavilov Institute of General Genetics, St. Petersburg Branch, Russian Academy of Sciences, 199034 St. Petersburg, Russia; Faculty of Science, University of Ostrava, Czech Republic, deceased; Laboratory of Amyloid Biology, St. Petersburg State University, 199034 St. Petersburg, Russia; Departments of Biochemistry and Molecular Biology; Microbiology and Pathology; Genetics Cell Biology and Anatomy, the University of Nebraska Medical Center, Omaha, NE, 68198, USA

## Abstract

In eukaryotes, translesion DNA synthesis (TLS) enables replication across DNA lesions, difficult DNA structures, and during replicative stress. While TLS prevents replication fork collapse, it is inherently mutagenic and requires tight regulation to prevent global mutation rates increase. Here, we show that elevated DNA polymerase pol ζ/Rev1 promotes spontaneous mutagenesis during replication.

The five subunits of pol ζ complex (pol ζ₅) are the catalytic subunit Rev3, the TLS polymerase Rev1, a dimer of accessory protein Rev7, the Pol31 and Pol32 subunits shared with replicative pol δ. Pol31 and Pol32 bind the Fe-S-containing C-terminal domain (CTD) of Rev3, which serves as a major assembly hub. We found that overproduction of pol ζ₅, but not four-subunit pol ζ₄ lacking Rev1 or Rev1 alone, increases spontaneous mutation rates by up to 20-fold, indicating that pol ζ₅ can access undamaged DNA during replication and compete with replicative DNA polymerases. This mutator effect is independent of PCNA ubiquitylation. augments by hydroxyurea and synergizes with DNA mismatch repair defects. Overproduction of a pol ζ₅ complex containing CTD-truncated Rev3 is also mutagenic, demonstrating that active TLS complexes can assemble and function without the canonical CTD interaction hub. Both complexes predominantly generate CG-to-GC transversions, a mutational pattern that resembles several COSMIC cancer mutational signatures. Inactivating Rev3 catalytic activity abolishes mutagenesis, whereas loss of Rev1 catalytic activity substantially reduces mutation rates and eliminates CG-to-GC transversions, identifying Rev1 as the primary cause of the observed mutation pattern. The findings reveal previously unrecognized mechanisms of pol ζ-dependent mutagenesis with broad implications for cellular evolution and cancer development.

## Introduction

Eukaryotic DNA replication balances the robustness of leading- and lagging-strand synthesis with fidelity. Highly efficient and accurate B-family DNA polymerases (pol) pol δ and pol ε, with help from primase/pol α in initiation, perform the two consecutive steps that ensure high accuracy of replication: nucleotide selectivity and exonucleolytic proofreading. The next step in assuring copied DNA quality is mismatch repair (MMR) (Kunkel and Erie 2015). The three steps working in series (Morrison et al. 1993), ensure high replication accuracy that, along with the other mechanisms, determine, according to most recent studies based on whole-genome sequencing, mutation rates that reach 10^-8^-10^-9^ per nucleotide per generation (Lynch et al 2008; Lujan et al. 2014; Lynch et al. 2023).

Threats to accurate replication include intrinsic and environmental factors that alter DNA structure. DNA damage blocks replicative polymerases because their snug active sites efficiently accommodate only correctly paired nucleotides (Kunkel and Bebenek 2000). Unless the cell repairs the damage or bypasses it via recombination, translesion synthesis (TLS) pols rescue replication through an elaborate mechanism that primarily uses Y-family pols (**Fig. 1A**) (Prakash et al. 2005). TLS pols possess a flexible active site that can accommodate the aberrant geometry of nascent base pairs (Yang and Gao 2018; Weaver et al. 2022). The low selectivity of TLS pols, favorable for bypass, makes TLS pols inaccurate while replicating intact DNA (Matsuda et al. 2000; Pavlov et al. 2006; McCulloch and Kunkel 2008). An additional problem of TLS is the extension of 3’ termini of the nascent DNA strand and damaged template that need specialized machinery. DNA pol ζ with REV1 and PCNA plays a central role in TLS in eukaryotes (Makarova and Burgers 2015; Martin and Wood 2019; Vaisman and Woodgate 2017). The complex’s primary roles are to recruit TLS machinery, extend mismatched primer termini after lesion bypass, and continue limited DNA synthesis before switching back to replicative polymerases. This process fixes mutations near damaged sites and can induce collateral mutations downstream of the lesion.(Prakash et al. 2005; Arianna and Korzhnev 2024; Kochenova et al. 2015; Póti et al. 2022)(**Fig. 1A**). Offsetting replication problems on damaged DNA improves viability at the cost of mutagenesis; blocking extension of aberrant termini prevents induced mutagenesis. Shifting the balance between replicative and TLS polymerases should elevate spontaneous mutagenesis (**Fig. 1B**), although DNA mismatch repair may correct many of these replication errors at active replication forks. .(Reyes et al. 2021).

**Figure 1.**
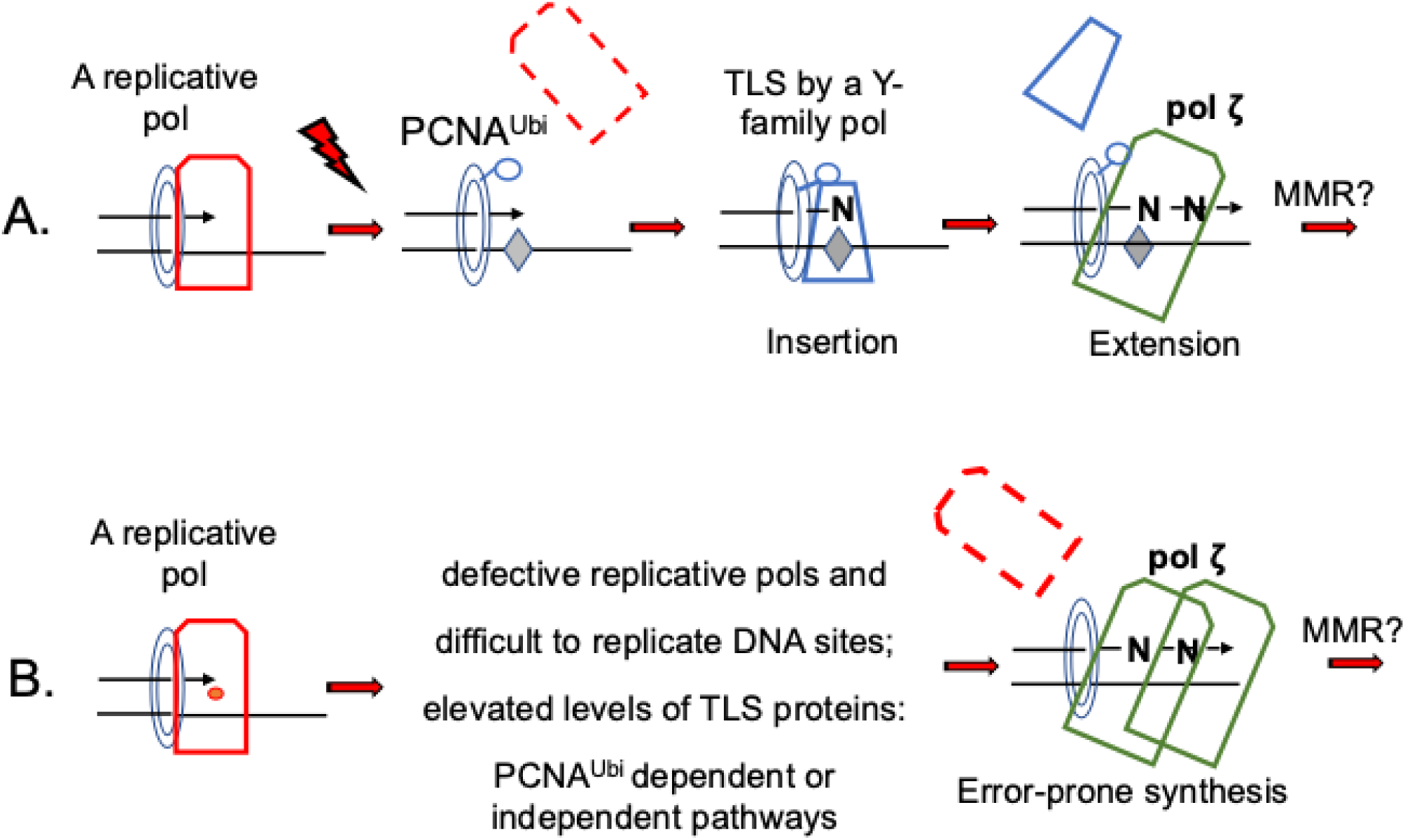
Role of pol ζ in translesion DNA synthesis (TLS). **A.** Processes and enzymes involved in TLS on damaged DNA and in induced/collateral mutagenesis. A grey diamond symbolizes a DNA damage site; the ring represents PCNA. The replicative polymerase is shown as a red chamfered rectangle; Y-family polymerases as a blue trapezoid; and pol ζ as a green chamfered rectangle. The letter “N” denotes any nucleotide, PCNA^UBi^ – monoubiquitylated PCNA, and MMR is DNA mismatch repair. **B.** Factors promoting the participation of pol ζ in replication of undamaged DNA, leading to spontaneous mutagenesis. The red dot in replicative pol represents an active-site alteration that reduces pol efficiency. The two green rectangles indicate increased pol ζ participation.

The low fidelity of TLS pols makes them dangerous for genome stability under normal conditions, requiring tight regulation. However, their ability to cause hypermutagenesis when regulatory control is lifted could be exploited when adaptation and evolution temporarily require high mutability. This is widespread in bacterial evolution, where up-regulated TLS (as a component of the SOS response) is crucial, dramatically elevating mutation rates in a hypermutable fraction of cells and conferring the adaptive potential (Radman et al. 1999; Maisnier-Patin and Roth 2015; Yamayoshi et al. 2018; Goodman 2016). In eukaryotes, transient, highly localized action of pol η, REV1, and pol ζ plays a seminal role in immunity, Including during diversification of lamprey antigen receptors and somatic hypermutation of immunoglobulin genes (Rogozin et al. 2001; Zeng et al. 2001; Krijger et al. 2013; Saribasak et al. 2012).

In budding yeast, no induced mutations occur when components of the TLS process and subunits of the specialized TLS four-subunit pol ζ in complex with REV1 (called pol ζ_5_, **Fig. 2A**) are absent or inactivated, for example, by changes in the Rev3 C-terminal domain (CTD) preventing Fe-S cluster binding (Lawrence 2004; Makarova and Burgers 2015) (**Fig. 2B**). Paradoxically, the complete deletion of the Rev3 CTD (**Fig. 2C**). is dispensable for ultraviolet radiation (UVR)-induced mutagenesis (Siebler et al. 2022; Siebler et al. 2014).

**Figure 2.**
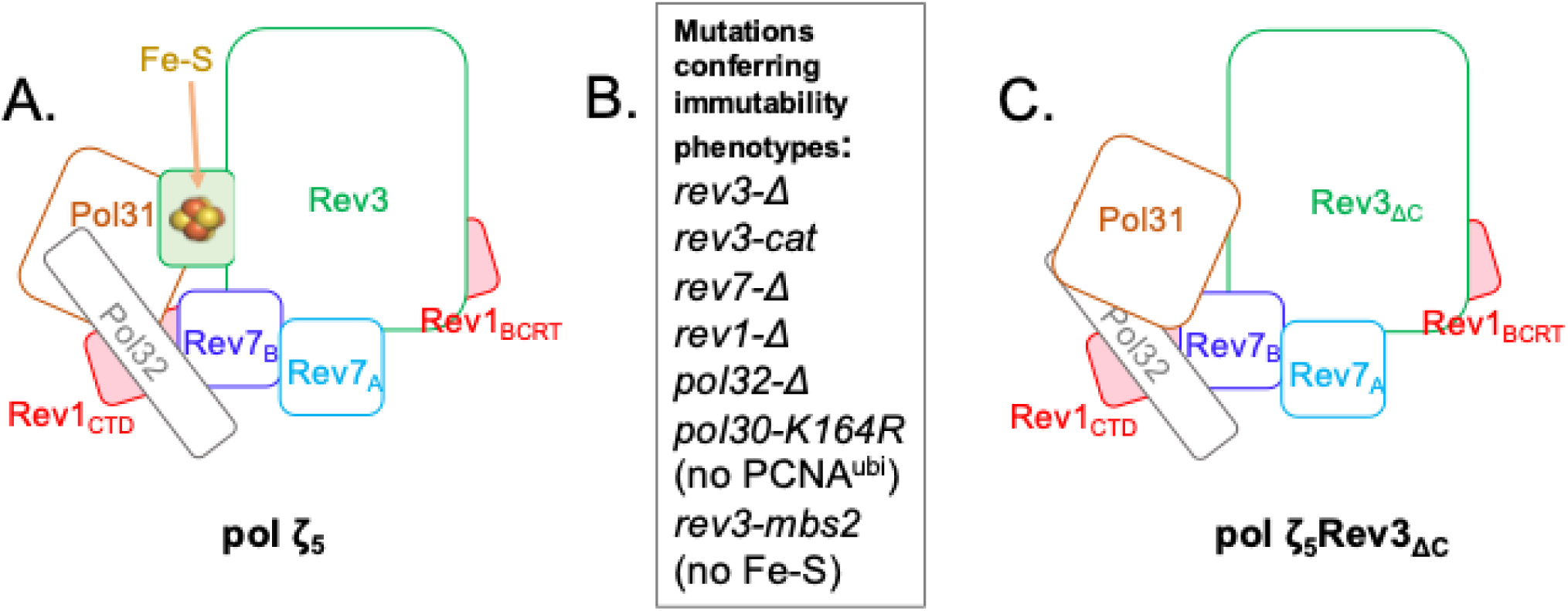
DNA polymerase ζ/Rev1 mutasome structure and function. Schematic structure of the 5-subunit yeast pol ζ and pol ζ variant unable to bind the Fe-S cluster. The CTD of the catalytic subunits with 4Fe-4S cluster (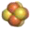) is shaded green. **B.** Mutations leading to the absence or severe reduction of induced mutagenesis. The index Δ in the allele name denotes a deletion. **C.** Schematic putative structure of the pol ζ-Rev3_ΔCTD_ variant without CTD.

Mutations inactivating REV3, the catalytic subunit of pol ζ, cause a two-fold antimutator effect in yeast (Quah et al. 1980) and human cells (Gyüre et al. 2023), suggesting that Pol ζ is responsible for around half of the spontaneous mutations. The involvement of pol ζ increases under replication stress caused by defects in replicative pols complexes (Pavlov et al. 2001b; Northam et al. 2006; Grabowska et al. 2014; Szwajczak et al. 2017), depletion of nucleotide pools (Northam et al. 2010), difficult-to-replicate DNA structures (Northam et al. 2014; Das et al. 2024; Ben Yamin et al. 2021), or elevated levels of TLS components (**Fig. 1B**). Pol ζ also participates in TLS necessary for completion of double-strand break recombination-repair, probable when processing damage in recessed single-stranded DNA and, therefore, introduces mutations during these transactions (Rattray et al. 2002; Malkova and Haber 2012). The role of TLS machinery in spontaneous mutagenesis in eukaryotes remains understudied. A few earlier reports examined whether raising TLS pols levels causes a mutator effect in eukaryotes (Pavlov et al. 2001a; Sasatani et al. 2017; Bebenek et al. 2001), but effects were marginal and sometimes indirect. A few recent studies suggest that TLS pols contribute to spontaneous mutations (Bainbridge and Daigaku 2024), but the genetic control and the mechanism remain unclear.

In the current study, we found that elevated levels of pol ζ_5_ (**Fig. 2A**) and its truncated variant, pol ζ_5_Rev3_ΔC_ (**Fig. 2C**), are mutagenic in yeast. We characterize this phenomenon in depth, determine mutation specificity, genetic control, and explore pol ζ structural requirements for efficient mutagenesis. The mutator effect is a valuable alternative readout of the functional state of pol ζ, beyond induced mutagenesis, helping to identify the minimal active mutasome and potentially explaining several CG-to-GC COSMIC mutation signatures of unknown etiology.

## Materials and Methods

### Plasmids for yeast strain construction

To generate a deletion of chromosomal *REV3,* we used one-step disruption by the *nat* gene, transforming yeast with PCR products on a pAG35 plasmid template and selecting for nourseothricin-resistant colonies (Goldstein and McCusker 1999). We verified the deletion by PCR with primers flanking *REV3.* To generate strains with *REV1* deletion disrupting the ORF with the ADE2 gene, we transformed appropriate strains by PCR products generated on the template chromosomal DNA of a yeast strain with an intact *ADE2* gene. To verify the disruption, we PCR-amplified a ∼4 kb *REV1*-encompassing region and distinguished between *REV1* and its ADE2 replacement by digesting with EcoRI. We list the primers needed for these manipulations in **sTable 1**.

To create a deletion of the chromosomal copy of the *REV7* gene, we used the integration-excision technique (Morrison et al. 1991). The DNA region flanking the ORF of *REV7* (**sFig. 1A**) was synthesized and cloned as a SalI-XbaI fragment into the pRS306 yeast vector by Azenta Life Sciences (Burlington, MA). We linearized the resulting plasmid with NcoI and used it to transform appropriate strains. We selected Ura+ transformants and plated them on FOA-containing medium to obtain clones that had lost the integrated plasmid. We identified deletion variants by PCR using primers for the *REV7* flanking region (**sTable 1**).

For correction of the *rad5* mutant allele, we used pRS306-*RAD5,* where *RAD5* was subcloned as a HindIII-XbaI fragment from pL2-1 vector (Fan et al. 1996) (gift from Dr. H. Klein laboratory, NYU).

To create PCNA mutants, we used one-step disruptions by SacI-cut pCH1572. plasmid with the mutation encoding for K164R change described in (Northam et al. 2006). To examine the effect of MMR deficiency, we disrupted the *MLH1* gene with *LEU2* using the *mlh1*Δ::*LEU2* plasmid described in (Prolla et al. 1994).

### Yeast strains

We created a series of yeast strains designed for studies of pol ζ *in vivo* and *in vitro* (**sFig. 2**, **sTable 2**). A founder strain, 1C-YPOM616 R5^+^, is a segregant of a hybrid from the cross of yJF1, used for purification of yeast replisome components (Yeeles et al. 2015) and LAN201, a descendant of CG379, used for studies of genetic effects of DNA pol mutations (Morrison et al. 1993; Bulock et al. 2020), and APOBEC deaminases (Lada et al. 2013). The main features of the strain are the deletion of the *PEP4* gene of the major protease (improves protein stability in extracts), deletion of the *BAR1* gene, barrier protein inactivating α-factor pheromone (providing a simple tool for synchronizing cells), and the presence of the wild-type *CAN1* gene (classic reporter to measure mutation rates and determine mutational specificity (Whelan et al. 1979)). We also corrected the Rad5 defect inherited from yJF1 by integrating and excising the WT *RAD5 gene* using a plasmid (Fan et al. 1996). Polymorphisms in DNA replication and repair genes between strains affect genome stability (Zhuk et al. 2023). Therefore, to obtain the 1C-YPOM616 R5^+^ complete genome (WT in the following text and figures), we re-sequenced its genome by combining Nanopore and Illumina technologies and used this verified genome to analyze gene expression by RNA-seq (see appropriate section). Next, we deleted the natural *REV3 and REV1* genes by gene disruption and *REV7* by integration-excision (**sFig. 1A**), either all three (strain 1C-YPOM616 R5^+^ Δr371) or individually, for complementation studies with wild-type and mutant alleles of the integrated genes (**sTable 2**). We integrated the constructs where five genes encoding pol ζ_5_ are expressed under a galactose promoter, similarly to the design for other replication factors (**Fig. 3AB**) (Yeeles et al. 2015), into 1C-YPOM616 R5^+^ rev3Δ (strains yζ_5v1_ and yζ_5_Rev3_ΔCv1_ that encode for wild-type Rev3 or for Rev3_ΔC_ without C-terminus), or 1C-YPOM616 R5^+^ Δr371, strains yζ_5v2_ and yζ_5_Rev3_ΔCv2_, respectively (**sTable 2**). For simplicity, we call the latter strains yζ_5 and and_ yζ_5_Rev3_ΔCTD_ in the manuscript’s text. We tested the functionality of *REV3rc, rev3rc-ΔC, REV7rc, and REV1rc* integrants by genetic complementation of appropriate single deletion strains (**sTable 3AB**).

**Figure 3.**
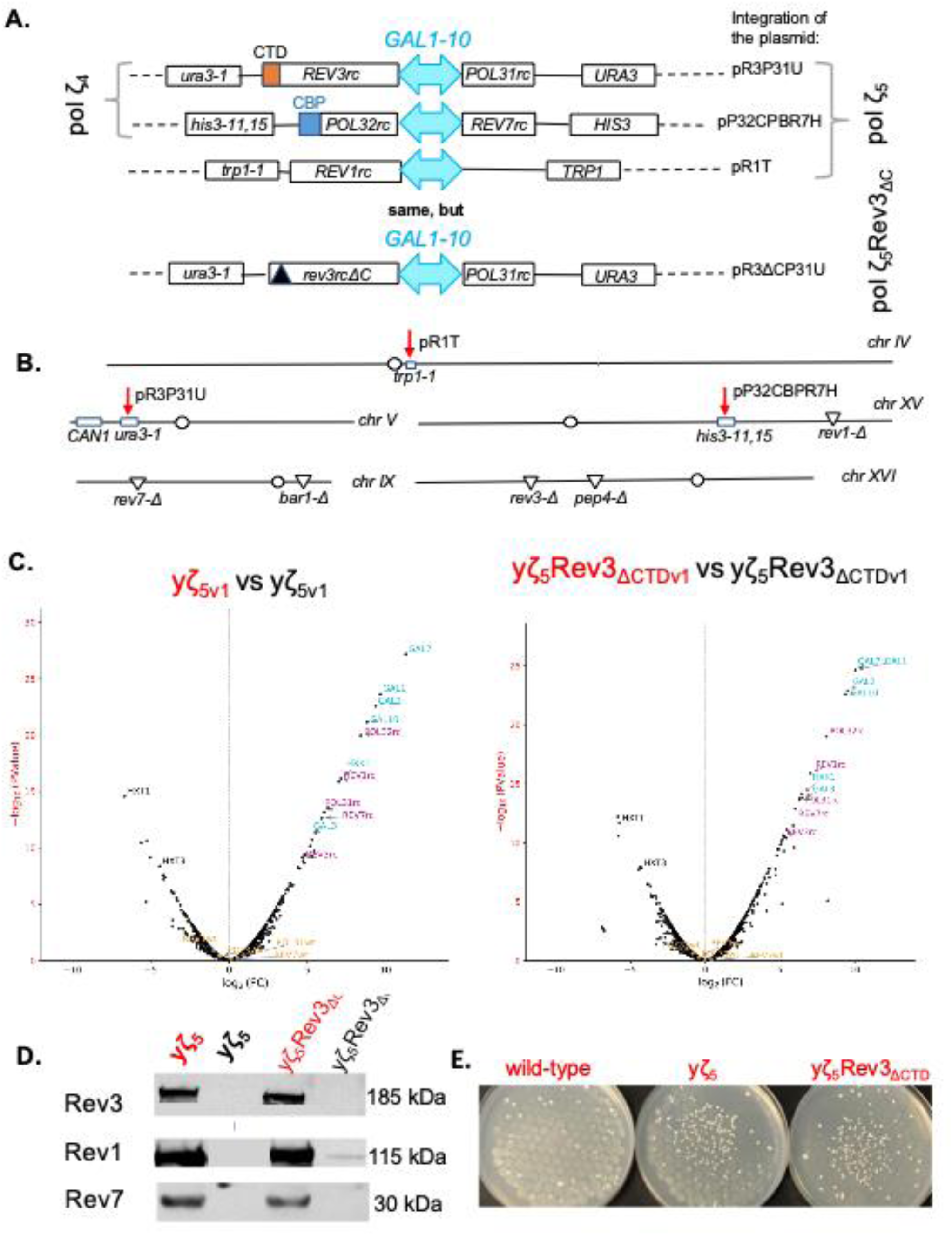
Basic features of strains overexpressing pol ζ in the current study. A. Basic features of strains overexpressing pol ζ. **A.** Changes in corresponding yeast chromosomes with deletion of natural genes but integrated genes for the expression of the four subunits of pol ζ and its associated pol Rev1. **B.** Part of the genome with integrated genes for the expression of the four subunits of pol ζ_4_ or five subunits of pol ζ_5._ or pol ζ-Rev3_ΔCTD._ **C.** Volcano plots of RNA-seq analysis of overexpression *(red) vs uninduced cultures (black) of yeast strains yζ_5v1._ or yζ_5_Rev3_ΔCTDv1_. **D.** Western blot of extracts with (red) or without (black) induction. **E.** Inducer (galactose) is spotted at the center of the plate and becomes a mutagen (a ring of mutant colonies).

### Media and growth conditions

We used standard yeast media, complete and minimal omission media (Rose et al. 1990). YPD was supplemented with adenine and uracil and named YPDAU and contained 2% glucose. We used YP_RAF_AU (2% raffinose instead of glucose) and YP_RAFGAL_AU (1% raffinose with 1% galactose) as needed. We grew yeast at 30 °C.

### Gene, oligonucleotide synthesis and site-directed mutagenesis

We purchased oligonucleotides for the one-step disruptions, PCR analysis of constructed strains, sequencing, and site-directed mutagenesis (**sTable 1**) from IDT (IA). Biomatik (Wilmington, DE, USA), custom-synthesized and cloned the recoded (codon-optimized, renamed *GENErc*) *REV3rc, rev3-ΔCrc, REV7rc,* and *REV1rc* genes into a bacterial vector, and we then re-cloned them into appropriate yeast expression vectors in our lab. The sequences of the re-coded genes are in **sFig. 1B**. Site-directed mutagenesis was performed according to (Zheng et al. 2004) with the use of Q5 DNA polymerase (NEB) or was performed by proprietary procedures by McLab (CA, USA) or BOC Sciences (NY, USA).

### Antibodies

AbClonal (MA, USA) produced custom polyclonal rabbit antibodies against parts of Rev3, Rev7, and Rev1 encoded by newly synthesized genes. Secondary anti-rabbit antibodies (ab97051) were from AbCam (MA).

### Genomic DNA sequencing and analysis

We streaked strains onto YPDAU agar plates (one strain per plate). We sent them to SeqCenter-LCC (PA) for genomic DNA isolation, combined Illumina and Nanopore sequencing, and preliminary genome assembly.

Base calling for Nanopore reads was performed with Guppy (v6.3.8; Oxford Nanopore Technologies), demultiplexing and adapter trimming for Illumina data were done with bcl-convert (v3.9.3; Illumina). Raw sequencing reads =quality was assessed by FastQC (v0.12.1, https://www.bioinformatics.babraham.ac.uk/projects/fastqc/), then aggregated with MultiQC (v1.28) (Ewels et al. 2016), with additional long-read quality assessment using NanoPlot (v1.44.1) (De Coster et al. 2018).

Illumina reads were trimmed with fastp (v0.24.1) (Chen et al. 2018), Nanopore reads were processed with Porechop (v0.2.4, https://github.com/rrwick/Porecho) and Chopper (v0.10.0) https://github.com/wdecoster/chopper) to remove adapters.

*De novo* genome assemblies were generated using Flye (v2.9.6) (Kolmogorov et al. 2019). Assemblies were polished with HyPo (v1.0.3) (Kundu r 2019) and scaffolded with RagTag (v2.1.0) (Alonge et al. 2022). Assembly completeness and quality were evaluated with BUSCO (v5.8.3) (Manni et al. 2021) and QUAST (v5.3.0) (Gurevich et al. 2013). To prevent mis-annotation of repetitive regions, genomes were soft-masked using RepeatModeler (v2.0.3) (Flynn et al. 2020) and RepeatMasker (v4.1.5, http://www.repeatmasker.org). Genome annotation was performed with GeMoMa (v1.9) (Keilwagen et al. 2019) and Augustus (v4.0.0) (Stanke et al. 2006), using protein-coding sequences and RNA-seq alignments as external evidence. The raw reads were deposited in NCBI sequence read archive (SRA), SUB16643933..

### RNA isolation, sequencing, and analysis

Yeast cultures were grown overnight in 3 mL of YP_RAF_AU. We then transferred 600 μl of each culture into 10 mL YPDAU and 800 μl into 10 mL YP_RAF_AU or YP_RAFGAL_AU to continue growing. After 4 hours, we pelleted the yeast cultures by centrifugation for 5 minutes at 2,500 rpm. Ιf RNA isolation was scheduled for the next day, we added RNA-Later (ThermoFisher, USA) and kept tubes at 4 °C. We then isolated RNA from the yeast cultures using the RiboPure™ - Yeast Kit (Ambion/Thermo Fisher Scientific, USA) according to the manufacturer’s directions. We measured the RNA concentration using a Denovix RNA Assay with a Qubit 2.0 Fluorometer (Invitrogen/Thermo Fisher Scientific). On average, RNA concentration was around 150 ng/μL. RNA samples were aliquoted and stored at -80 °C. We sent samples on dry ice to SeqCenter-LCC (PA, USA) for sequencing and preliminary bioinformatics analysis. We adjusted the sample concentration and volume per the service provider’s guidelines.

Sequencing was done on a NovaSeq 6000, generating 2×51 bp reads. Demultiplexing, quality control, and adapter trimming were performed with bcl-convert (v4.0.3).

Raw reads were evaluated with FastQC (v0.12.1) and MultiQC (v1.28) as described for DNA reads above. followed by trimming with fastp (v0.24.1) (Chen et al. 2018). rRNA was removed using bbduk from the BBTools package https://www.osti.gov/servlets/purl/1241166, with rRNA sequences extracted from the S288C reference genome (GCF_000146045.2).

RNA-seq reads were aligned to the final genome assemblies with STAR (Dobin et al. 2013). Read counts were generated with featureCounts (from the Subread package) (Liao et al. 2014). Differential gene expression analysis was carried out with edgeR (Robinson et al. 2010); significance of expression changes and pathway enrichment were assessed, and results are presented as volcano plots.

The data are presented as volcano plots. Raw RNAseq reads will be deposited as above. The pipeline, including genome assembly, annotation, and all RNA-seq processing steps, is available at https://github.com/mattpanteleev/GAD-pipeline.

### Yeast protein extracts and Immunoblots

Western blots were used to show that the mutants studied did not affect protein stability in extracts. Logarithmic yeast cultures grown in either 10 mL of YP_RAF_AU or YP_RAFGAL_AU (4 hours of induction) were pelleted by centrifugation, washed in PK lysis buffer (50 mM Tris-Cl pH7.6, 50 mM NaCl, 0.1% Triton X-100, 0.1% Tween 20, 1 mM EDTA), (Pessoa-Brandao and Sclafani 2004). Cells were disrupted in a VortexGenie 2 equipped with a multi-tube horizontal holder (two times for 2 min at max speed alternated by 1 min chill on ice) with 500 μl cold glass beads in 500 μl PK with 0.5 mM PMSF. We noticed that adding 1x EDTA-free Halt Protease inhibitor (Thermo Scientific) helped prevent extensive protein degradation. Cell debris and beads were pelleted by centrifugation at 14,000g for 15 min at 4 °C, and the supernatant was transferred to a new tube. We determined total protein concentration by the Bradford method. The method reproducibly yields extracts with around 10 ng/μL of total protein. The appropriately diluted extract was mixed with loading buffer, denatured at 80 °C for two minutes, and loaded (60 μg of total protein) into wells of 4-12% or 10% NuPage Bis-Tris gels. To estimate protein sizes, we used a PageRuler Plus Prestained Protein Ladder covering 10 to 250 kDa (Thermo Fisher Scientific). After 40 min electrophoresis at 200V in appropriate buffer (MOPS or MES), proteins were transferred to a PVDF membrane (Thermo Fisher Scientific) in an XCell Blot Module (Invitrogen) at 30 V for 1 hour. The membrane was washed 2 x 2 min in water, then blocked in 14 mL of TBS Starting Block for 1 h at room temperature with agitation. The membrane was then incubated with appropriate primary antibodies (or a combination) at a 1:500 dilution in 14 mL of TBS. Starting Block (either 1 hour at room temperature or overnight at 4 °C with gentle rocking). Then, after washing twice for 5 min with 15 mL TBST followed by 3 times 5 min with 15 mL TBS, the membranes were incubated at room temperature for 1 h in 14 mL of TBS Starting Block with the horseradish peroxidase-conjugated goat anti-rabbit IgG secondary antibodies diluted at 1:3500. Membranes were washed four times for 5 min with 15 mL TBST followed by four times 5 min with 15 mL TBS. We detected proteins using either a chemiluminescence kit (Pierce™ ECL Western Blotting Substrate, Thermo Fisher) with Thermo Fisher Scientific CL-XPosure film or an Ultra-TMB blotting solution (Pierce/Thermo Fisher) for chromogenic detection.

### Spot tests, mutation rates, and mutant frequencies determination

For a galactose-induced mutation spot test, cells grown overnight in YP_RAF_AU were plated on the same medium, with 20 μl of 20% galactose spotted in the center of the plates and dried. After 1-day incubation, the growing cells were replica-plated onto synthetic complete medium containing canavanine. We recorded resistant colonies after 4 days of further growth.

Cadmium-sensitivity spot tests were done similarly by spotting 10-15 μL of 50 mM CdCl2 solution in the center of raffinose, galactose, or YPDAU plates, waiting for the spot to dry, and applying fresh yeast cultures with a multi-prong device. After 24 hours of incubation, we replica-plated cells onto the selective medium, as above.

For quantitative mutation analysis, we grew individual colonies for 2-4 hours in YP_RAF_AU. The culture was split 1:1 and added to equal amounts of fresh medium with 2% raffinose in one tube (uninduced control) or 2% galactose in the other (induction of genes under *GAL* promoters). The rate of *CAN1* forward mutation (the reporter detects all types of mutations, including base pair substitutions, frameshifts, large deletions, and chromosome arm loss) was measured by fluctuation analysis as described (Shcherbakova and Kunkel 1999; Pavlov et al. 2001b). The median rate was calculated by Drake’s formula (Drake 1991), as described before, from at least nine clones of the same strain for each experiment (repeated three times). UV light mutagenesis was studied as before (Siebler et al. 2014; Stepchenkova et al. 2017). We plated appropriate cell dilutions on selective and complete plates and irradiated them. We calculated the mutant frequency in each culture as the ratio of mutants on selective plates to surviving, colony-forming cells on complete plates, multiplied by the dilution factor. We determined mutant frequency at all UV doses for 2-3 cultures of the same strain. We calculated the UV-induced mutant frequency by subtracting the spontaneous background frequency from the mutant frequency in UV-irradiated cells. We treated cells with hydroxyurea (HU) as described in (Northam et al. 2010), but at lower concentrations, and grew HU-treated cells overnight in either YP_RAF_AU and YP_RAFGAL_AU.

### Analysis of mutation specificity

The *CAN1* locus from independent Can^R^ colonies was amplified and sequenced as described (Siebler et al. 2022). We also used bulk analysis by NGS as described (Jiang et al. 2021; Jiang et al. 2022). We aligned sequences to the wild-type *CAN1* allele sequence, with polymorphisms characteristic of the LAN201 strain, and detected and mapped mutations to form spectra. Detailed procedures for mutation-spectrum analysis and comparisons to cancer mutation signatures are described in **sFile. 1**.

## Results

### Properties of yeast strains with regulatable expression of pol ζ_5_

When the five genes encoding pol ζ_5_ or a variant for CTD of Rev3 truncation, pol ζ_5_Rev3_ΔCTD_ (**Fig. 2A,C**), are expressed under galactose promoter, which can be regulated by various amount of galactose like rheostat, the relative gene expression profiles were similar in yζ_5_ and yζ_5_Rev3_ΔC_ (**Fig. 3C**). In strains when only one natural gene, *REV3,* was deleted (yζ_5v1_ and yζ_5_Rev3_ΔCv1_, **sTable 2**) we can directly compare mRNA production of galactose-inducible genes with their natural counterparts (**Fig.2C**, **sFig. 3**). On average, induction caused a substantial increase in mRNA levels, and the two strains behaved identically. We conclude that the allele truncating the Rev3 CTD did not affect RNA production or stability. Only a few genes besides galactose-regulated ones were induced or inhibited by galactose (**Fig. 3C**). Next, we examined protein production in strains with newly constructed optimized *REV3*, REV7, and *REV1* genes by Western blot and found that Rev3. Rev7 and Rev1 were robustly detected only in cell extracts of yeast grown in galactose (induction conditions), consistent with RNA-seq data (**Fig. 3D**). Deletion of the Rev3 CTD did not affect the stability of the proteins studied.

Elevated production of pol ζ_5_ or polζ_5_Rev3_ΔC_ did not affect yeast growth but robustly induced *can1* mutants by galactose in spot tests (**Fig. 3E**). Expression of pol ζ_4_ and Rev1 alone had no mutator effect (**sTable 3**). We conclude that expressing the five genes encoding pol ζ is sufficient to assemble a minimal mutasome that participates in spontaneous mutagenesis. Further work explores the effect in detail.

### Parameters and genetic control of mutator effects in strains with elevated pol ζ levels

Galactose became a novel mutagen in our strains because it induced pol ζ/Rev1 expression, producing an active “mutasome” (**Fig. 3E**). We compared spontaneous and UVR-induced mutagenesis in the parent strain with natural pol ζ genes (WT), yζ_5_ and yζ_5_Rev3_ΔC_ strains and their variants (**Fig. 4**, **Fig. 5, sTables 3 and 4**). Without induction, strain yζ_5_ is like WT: we recorded the same level of spontaneous and UVR-induced mutations. Increased pol ζ_5_ expression increased spontaneous mutation rates10-20-fold but did not change UVR mutagenesis levels. When the four cysteines in MBS2 of Rev3 are substituted with alanines, UVR mutagenesis without galactose induction is ablated, as documented earlier (Baranovskiy et al. 2012); however, after induction, both spontaneous and UV-induced mutagenesis reach levels like those in strains with wild-type *REV3.* (**sTable 4).** The yζ_5_Rev3_ΔC_ strain showed slightly lower UVR-induced mutagenesis under low expression, consistent with published data for the same deletion in natural gene (Siebler et al. 2014). Both mutagenesis phenotypes reached the yζ_5_ level at high expression. We conclude that elevated expression generated enough active pol ζ_5_ complexes to restore WT UVR-induced mutagenesis and produce a robust spontaneous mutator phenotype, even when CTD was unable to bind the Fe-S cluster or was completely absent.

**Figure 4.**
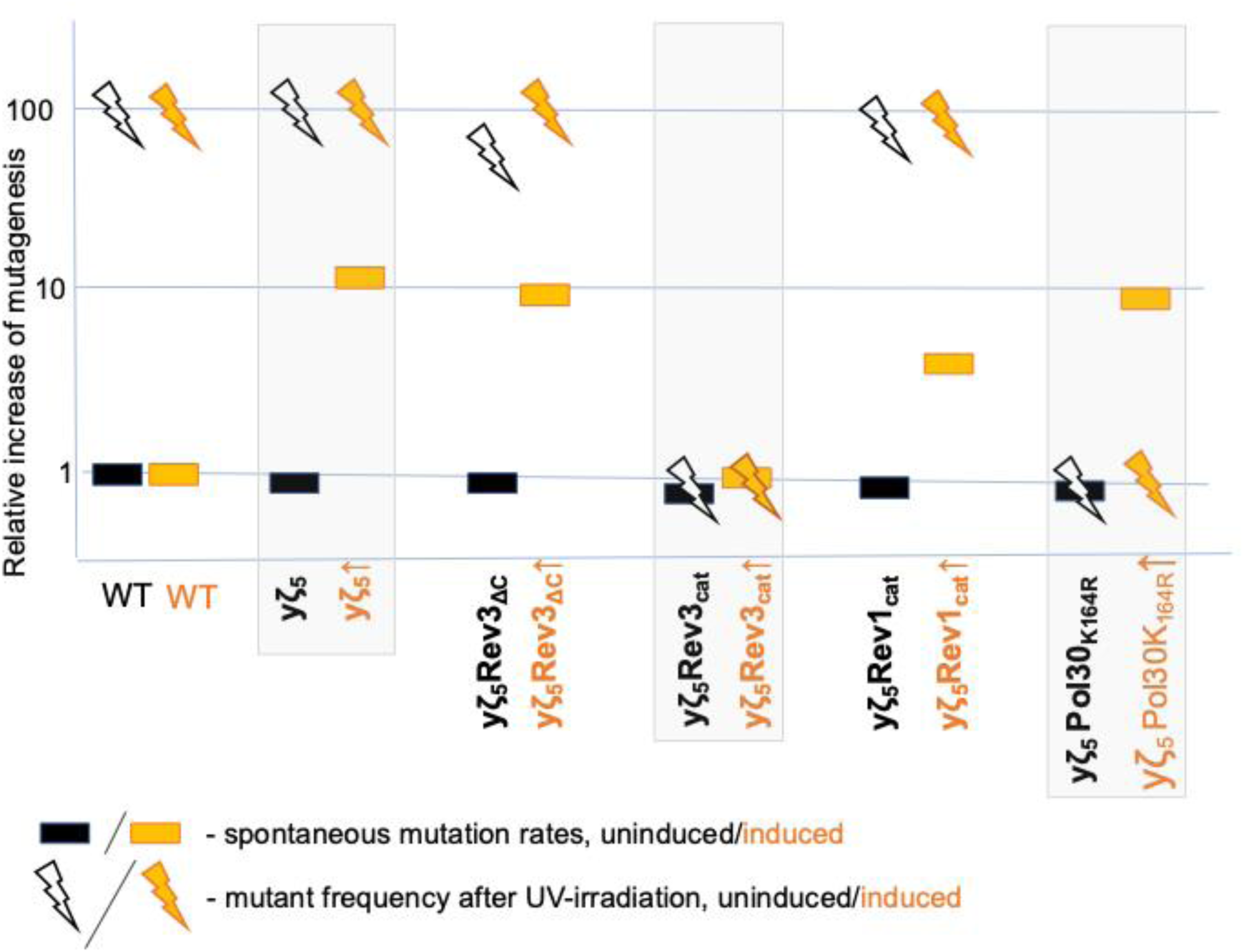
Summary of genetic control of spontaneous and ultraviolet light irradiation (UVR)-induced mutagenesis in strains with different expression levels of pol ζ genes and alleles. Elevated pol ζ levels increase spontaneous mutation rates, and this effect depends on the catalytic activities of the polymerases Rev3 and Rev1 while only Rev3 catalytic activity is absolutely required for UVR mutagenesis. The increase of spontaneous mutagenesis is PCNA-ubiquitylation-independent, in contrast with UVR mutagenesis. The spontaneous mutation rate (shown in rectangles) of Can^r^ mutations in the WT strain (3×10^-7^) is set to 1, and UVR elevates mutation frequency (shown as lightning) 100-fold. We compare the effects of mutations in yζ_5_ strains with (yellow) or without induction (black). The variants are yζ_5_ strains with alleles encoding deletion of the the CTD of Rev3 (Rev3_ΔC_), catalytically dead mutants of Rev3 and Rev1 (Rev3_cat_ or Rev1_cat_), and yζ_5_ strains in which PCNA cannot be ubiquitylated (Pol30K164R).

**Figure 5.**
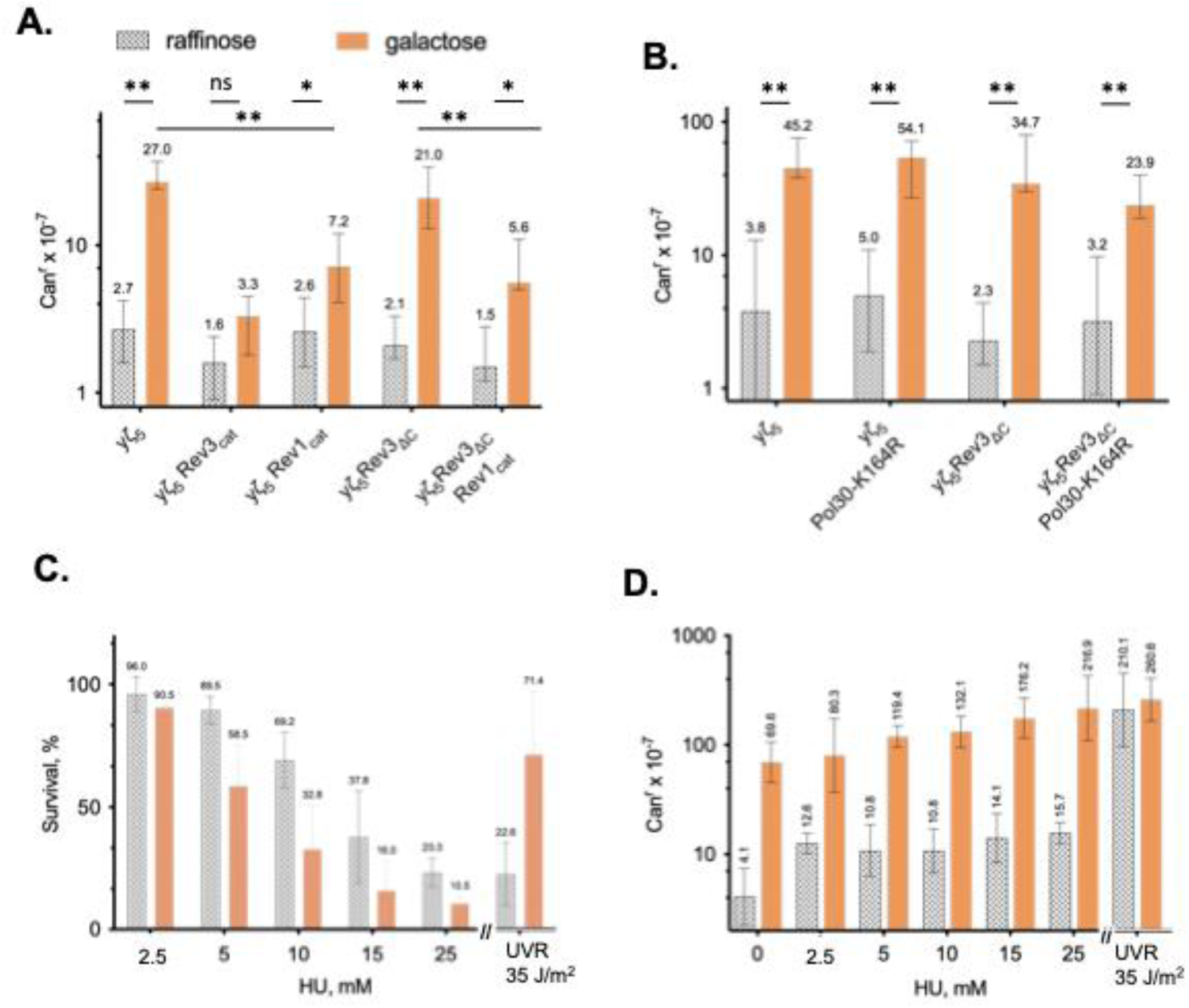
Spontaneous mutagenesis by Pol ζ requires Rev3 catalytic activity, depends only partially on Rev1. (A), is independent of PCNA ubiquitylation (B), and is multiplicative with the mutagenic effect of HU (C, survival, D mutagenesis, strain yζ_5_). Statistical differences for A and B are labeled: ns – not significant, * P<0.05, ** P<0.01; Wilcoxon-Mann Whitney test.

Both UVR-induced and elevated spontaneous mutagenesis are abrogated when Rev3 is catalytically dead (**Fig. 4**, **Fig. 5A**). In contrast, catalytically dead Rev1 did not affect UVR, consistent with the literature (Wiltrout and Walker 2011), but substantially (∼ 3-fold) lowered spontaneous mutagenesis. Deletion of the M1 region, which was shown to elevate Rev1 catalytic activity (Makarova et al. 2014), did not affect spontaneous or induced mutation rates/frequencies, with or without induction (**sTable 4**), consistent with the literature (Bezalel-Buch et al. 2026). Elevated *RAD30* expression did not affect mutagenesis in strains with elevated levels of pol ζ, as previously shown in WT strains (Pavlov et al. 2001a). UVR-induced mutagenesis in the same strains, including strains with the *pol30-K164R* mutation, where PCNA cannot be monoubiquitylated (Stelter and Ulrich 2003), is severely suppressed (**Fig. 4**; results are shown for the y ζ_5_ strain; truncated Rev3 behaves similarly). Unexpectedly, the spontaneous mutator effect of the expression of both forms of pol ζ was independent of PCNA^Ubi^ (**Fig. 4**, **Fig. 5B, sFig. 4A**). Mutagenesis by defective replisome causing the recruitment of pol ζ to replication of undamaged DNA is PCNA^ubi^-dependent (Northam et al. 2006). Therefore, yeast expressing pol ζ exploits a special, novel mechanism of its participation in replication.

The inhibition of MMR by Cd^2+^ (Jin et al. 2003) synergized mutagenesis by elevated pol ζ as it does for canonical mutator replicative pols (**sFig. 4B**, positive controls were strains with proofreading-defective replicative pol ε (*pol2-4* (Morrison et al. 1991), a cancer-associated variant *pol2-V411L* (Barbari et al. 2022)), or pol δ (mutation *pol3-5DV*, (Jin et al. 2005)), arguing for a direct participation of pol ζ in S-phase replication under our conditions. To further access the role of MMR, we generated MMR-defective strains that overexpress pol ζ and Rev1. We found that the elevated levels of pol ζ_5_ synergize with disruption of one of the key MMR genes*, MLH1* **Table 1**. Each factor alone elevated mutation rates 10-20-fold, but combining an MMR defect with elevated pol ζ caused a 65-126-fold increase. The synergy value falls between additivity and multiplicity, meaning that different error types leading to mutations in the *CAN1* gene are not corrected with equal efficiency. We observed that MMR deficiency has slightly (but not significantly, P=0.108) higher in yζ_5_Rev3_ΔC_ than in yζ_5_ strains. This tendency might indicate differences in MMR efficiency for errors made by these two complexes. We conclude that MMR corrects a substantial proportion of pol ζ errors, suggesting that the polymerase participates in S-phase replication. In this respect, the spontaneous mutator effect does not mimic the parameters of Rev3-dependent mutagenesis in strains with defects of pol ε subunits Dpb3 and 4 (Aksenova et al. 2010), but is like Rev3-dependent mutagenesis in strains with defects in replicative DNA polymerases (Pavlov et al. 2001b). Further elucidation of the mechanism is necessary because TLS polymerases are recruited to replication of undamaged DNA in human cells under various stress conditions, for example, hypoxia (Yehuda et al. 2025) or pathology (Anand et al. 2023). To this end, we examined the effects of hydroxyurea, which inhibits ribonucleotide reductase, depletes dNTP pools, and thus slows replication. Previously, HU was shown to be moderately mutagenic in yeast, and this mutator effect is pol ζ-dependent (Northam et al. 2010), implying pol ζ participation in replication under stress. Because pol ζ, unlike major replicative DNA pols, is tolerant to low dNTP concentrations (Kochenova et al. 2017), we expected that HU would augment pol ζ mutagenic effects. Indeed, HU increased Can^r^ mutation frequency 3-fold at 25 mM and reduced the number of viable cells; increasing pol ζ levels caused a 17-fold increase (**Fig. 5C and D**). Together, these factors led to a multiplicative 53-fold increase in mutation rates and a further decrease in cell viability. Interestingly, while pol ζ expression rescued the lethal effects of UV, it potentiated HU-induced cell lethality (**Fig. 5C**).

**Table 1.** MMR removes errors made during elevated production of pol ζ.

| Strain number | Principal genotype | Presence of the inducer, galactose | Median, rate 10<sup>-7</sup> (95% CI) | 95% CI | Fold increase above wild-type |
|---|---|---|---|---|---|
| yo2595 | wt | + | 1.8 | 1.2-5.1 | 1 |
| yo3480 | *mlh1Δ::LEU2* | + | 26.2 | 24.7-57.2 | 14.5 |
| yo3111 | genes for pol ζ<sub>5</sub> | - | 0.8 | 0.7-1.9 | 0.4 |
|  |  | + | 29.3 | 21.6-75.2 | 16.2 |
| yo3471 | genes for pol ζ<sub>5</sub><br>*mlh1Δ::LEU2* | - | 33.2 | 23.6-83.3 | 18.4 |
|  |  | + | 117 | 52-194 | 65 |
| yo3118 | pol ζ<sub>5</sub> Rev3<sub>ΔC</sub> | - | 1.1 | 0.7-1.5 | 0.6 |
|  |  | + | 43.5 | 30.0-77.4 | 24 |
| yo3474 | genes for pol ζ<sub>5</sub> Rev3<sub>ΔC</sub><br>*mlh1Δ::LEU2* | - | 29.1 | 19.4-48.2 | 16.1 |
|  |  | + | 228 | 103-403 | 126 |

### Differential effects of amino acid substitutions affecting interactions in pol ζ complex: yζ_5_ versus yζ_5_Rev3_ΔC._

We examined the roles of accessory subunits in pol ζ assembly. The possibility of defining critical elements necessary for the complex’s function comes from the finding that induced and spontaneous mutasome functions of pol ζ withstand the loss of the CTD hub of the catalytic Rev3 subunit. Interactions via Rev7/Rev1, or novel interactions between Rev3ΔC and Pol31, appear critical (schematically illustrated in **Fig. 2A** versus **C**). We changed several amino acids in Rev7 that were predicted to be significant for inter-subunit interactions by structural and biochemical studies (Hara et al. 2010; McPherson et al. 2021; Acharya et al. 2005; Wojtaszek et al. 2012; Kikuchi et al. 2012; Pozhidaeva et al. 2012; Pustovalova et al. 2016; Malik et al. 2020; McPherson et al. 2023; Malik et al. 2024). **Table 2AB and sTable 4** summarize the results. Overall, these changes had the most profound effects in yζ₅Rev3ΔC strains, providing strong evidence that Rev7 serves as a critical hub for assembly of functional pol ζ₅ complexes and that even a single amino acid substitution can drastically impair complex function.

**Table 2.**
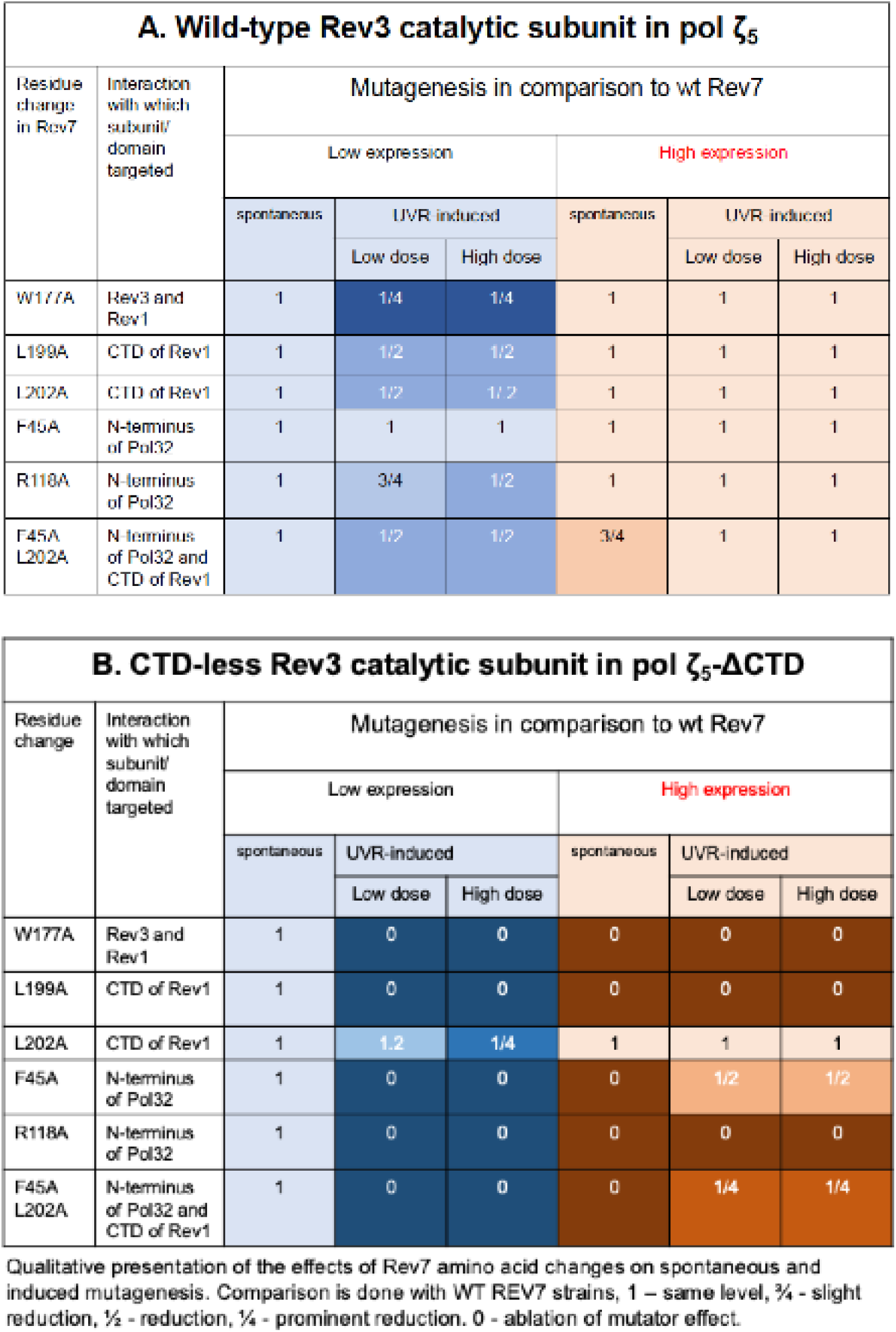
The effect of Rev7 mutations depends on the status of Rev3.

| **A. Wild-type Rev3 catalytic subunit in pol ζ<sub>5</sub>** |  |  |  |  |  |  |  |
|---|---|---|---|---|---|---|---|
| Residue change in Rev7 | Interaction with which subunit/ domain targeted | Mutagenesis in comparison to wt Rev7 |  |  |  |  |  |
|  |  | Low expression |  |  | High expression |  |  |
|  |  | spontaneous | UVR-induced |  | spontaneous | UVR-induced |  |
|  |  |  | Low dose | High dose |  | Low dose | High dose |
| W177A | Rev3 and Rev1 | 1 | 1/4 | 1/4 | 1 | 1 | 1 |
| L199A | CTD of Rev1 | 1 | 1/2 | 1/2 | 1 | 1 | 1 |
| L202A | CTD of Rev1 | 1 | 1/2 | 1/ 2 | 1 | 1 | 1 |
| F45A | N-terminus of Pol32 | 1 | 1 | 1 | 1 | 1 | 1 |
| R118A | N-terminus of Pol32 | 1 | 3/4 | 1/2 | 1 | 1 | 1 |
| F45A<br>L202A | N-terminus of Pol32 and CTD of Rev1 | 1 | 1/2 | 1/2 | 3/4 | 1 | 1 |
| **B. CTD-less Rev3 catalytic subunit in pol ζ<sub>5</sub>-ΔCTD** |  |  |  |  |  |  |  |
| Residue change | Interaction with which subunit/ domain targeted | Mutagenesis in comparison to wt Rev7 |  |  |  |  |  |
|  |  | Low expression |  |  | High expression |  |  |
|  |  | spontaneous | UVR-induced |  | spontaneous | UVR-induced |  |
|  |  |  | Low dose | High dose |  | Low dose | High dose |
| W177A | Rev3 and Rev1 | 1 | 0 | 0 | 0 | 0 | 0 |
| L199A | CTD of Rev1 | 1 | 0 | 0 | 0 | 0 | 0 |
| L202A | CTD of Rev1 | 1 | 1.2 | 1/4 | 1 | 1 | 1 |
| F45A | N-terminus of Pol32 | 1 | 0 | 0 | 0 | 1/2 | 1/2 |
| R118A | N-terminus of Pol32 | 1 | 0 | 0 | 0 | 0 | 0 |
| F45A<br>L202A | N-terminus of Pol32 and CTD of Rev1 | 1 | 0 | 0 | 0 | 1/4 | 1/4 |

Qualitative presentation of the effects of Rev7 amino acid changes on spontaneous and induced mutagenesis. Comparison is done with WT REV7 strains, 1 – same level, ¾ - slight reduction, ½ - reduction, ¼ - prominent reduction. 0 - ablation of mutator effect.

### Prominent role of Rev1 in establishing mutation spectra of pol ζ

We sequenced hundreds of Can^r^ mutants arising when both variants, pol ζ_5_ or pol ζ Rev3_ΔCTD,_ are elevated and presented the acquired information on types of mutations (**sTable 5A,** pie graphs, **Fig. 6A**), types and positions on traditional mutation spectra (s**Fig. 5**). Note that the *CAN1* spectra approximate the genomic mutation spectra (Lynch et al. 2008). The mutation spectra of yζ₅ and yζ₅Rev3ΔC were similar at the stringent 0.01 significance level (**sTable 5B**), providing further evidence that pol ζ containing the Rev3 CTD truncation functions similarly to the wild-type mutasome. GC→CG transversions dominated the spectra, accounting for more than 50% of all mutations. These spectra are very different from spectra of spontaneous mutation in the *CAN1* gene (**sTable 5B, Fig. 6A**, leftist pie, data for 537 wild-type *can1* mutants in related genetic backgrounds are from (Ohnishi et al. 2004; Lang and Murray 2008; Aksenova et al. 2010; Grabowska et al. 2014; Garbacz et al. 2015)). Mutation hotspots also frequently (in 31 sites) overlapped (**Fig. 6B, sFig. 5).** Since both UVR and spontaneous mutagenesis are abrogated when Rev3 is catalytically dead (**Figs. 4 and 5A**), we did not analyze the nature of Can^R^ mutants in these variants. Catalytically dead Rev1 substantially lowered spontaneous mutagenesis. Most GC-to-CG transversions vanished from the mutation spectrum (**Fig. 6A**), suggesting that most mutations in pol ζ_5_-expressing strains arise from misincorporation of dCTP by Rev1’s transferase activity.

**Figure 6.**
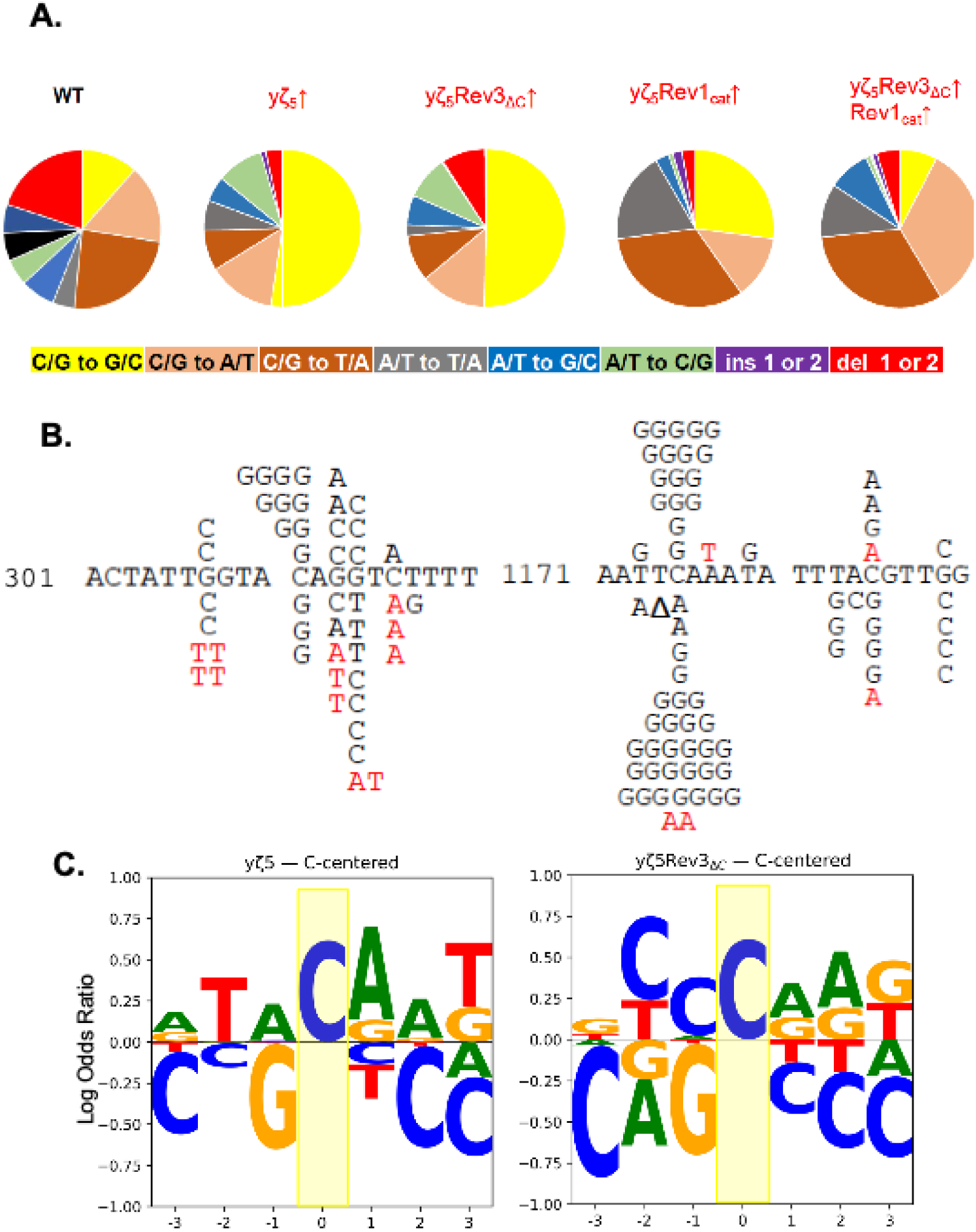
Mutation specificity of mutasome. **A.** Prominent role of Rev1 in driving GC to CG transversions. **B.** Examples of pol ζ variants’ mutational hotspots in the *CAN1* gene. Black letters/symbols: changes found in yζ_5_ strains are above the wild-type DNA sequence, and changes in yζ_5_Rev3_ΔC_ are under it. Changes in corresponding Rev1cat strains are in red. Δ stands for a one-nucleotide deletion. **C.** Subtle alterations in the specificity of spontaneous mutation in strains with CTD-less pol ζ (LOGOs).

Interestingly, several hotspots for pol ζ mutagenesis remained in the spectra with Rev1cat. However, the types of mutations changed, e.g, at positions 313, 895, 937, 1163, 1211 (**sFig. 5**). Collapsing substitution classes (**sTable 5B)** revealed that the major source of spectra divergence between strains is the proportion of Rev1-dependent mutations (C>G), which is strongly elevated in yζ_5_ and yζ_5_Rev3_ΔC_ strains and almost completely abolished by Rev1 catalytic inactivation. Note that spontaneous G to C mutations in wild-type strains are rare (**sTable 5B**). Because Rev1 expression alone did not increase spontaneous mutations, we conclude that Rev1 interacts with both forms of pol ζ during replication of undamaged DNA. However, the interaction differs, as evidenced by the switch in abundance from CG to GC and from GC to TA in pol ζ_5_Rev1_cat_ vs pol ζ_5_Rev3_ΔC_Rev1_cat,_ and by the increased proportion of small deletions in strains with truncated Rev3 (**sTable 5A**). The analysis of context specificity of G to C mutations (**Suppl. File 1**) also revealed differences between yζ_5_ and yζ_5_Rev3_ΔC_ strains (**Fig. 6C**). Interestingly, in both cases, template purines at positions +1 and +2 relative to the mutation site were preferred over pyrimidines (the sequence shown is replicated from right to left).

### Mutation signatures of elevated pol ζ mirror mutation SSBs in cancers

The mutation distributions in strains expressing different pol ζ variants versus wild-type were quite different, with much more uniformity in the latter (**sFig. 6**), suggesting different mechanisms of mutation occurrence. In the wild-type strain, multiple factors contribute to mutation induction, including replicative DNA pol errors, TLS processing of spontaneous DNA damage, and DNA structures that cause mutations when mutasome outcompetes replicative pols at sites where they are less efficient. We found that the spectra resemble several mutational signatures in cancer (**Fig. 7**, s**Table 6, sFig. 7**). The most prominent is similarity to signature SBS39, of unknown etiology. The moderate correlation suggests that pol ζ contributes to certain cancers, either through inadvertent elevated expression or through untargeted mutagenesis.

**Fig 7.**
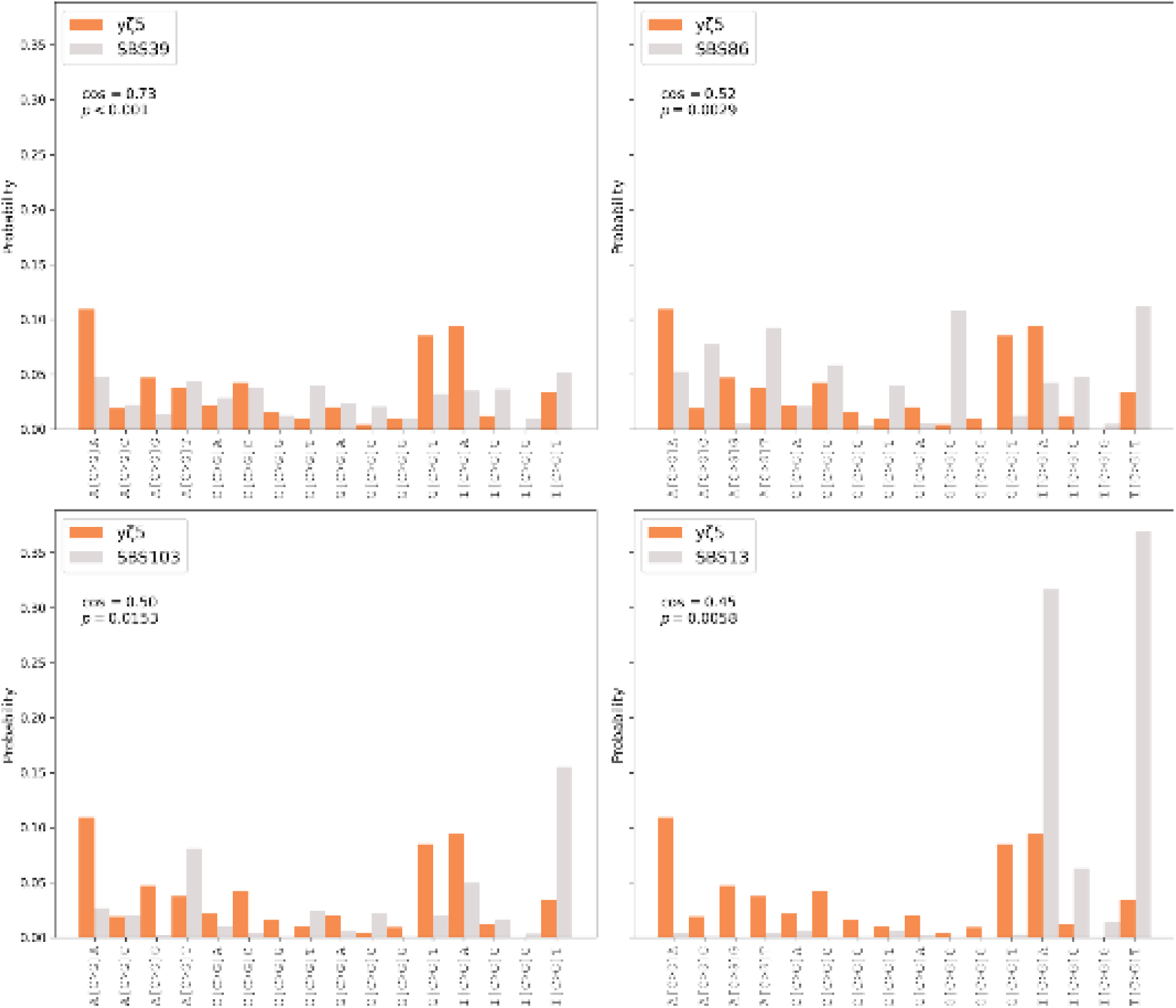
DNA context specificity of G to C transversions caused by elevated production of pol ζ coincides with cancer mutation signatures. Cosmic mutation signatures are shown as gray bars; brown-orange bars show the pol ζ_5_ signature.

## Discussion

We found that elevated levels of pol ζ₅ exert a spontaneous mutator effect in yeast, fulfilling the prediction shown in **Fig. 1B**. This mutagenesis occurs independently of PCNA ubiquitylation and is exacerbated by defects in mismatch repair (MMR). We defined the minimal composition of the mutasome responsible for this phenotype and determined the extent to which the requirements for spontaneous and induced mutagenesis overlap. One of the most significant findings was the prominent contribution of Rev1 to GC→CG transversions. Notably, the sequence context of these mutations closely resembles several COSMIC SBS signatures of unknown etiology, suggesting that analogous mutagenic mechanisms may underlie mutation accumulation in certain human cancers.

### Mechanism of mutator effect by elevated production of pol ζ_5_

Defects in pol ζ in the absence or with non-functional Rev3 reduce spontaneous mutagenesis in yeast two-fold, implying that half of mutations originate from the activity of this specialized polymerase (Quah et al. 1980; Abdulovic et al. 2008; Szwajczak et al. 2017). The involvement is substantially elevated when replication is impeded by replication stress caused by impairment of replicative DNA polymerase complexes (Northam et al. 2006; Aksenova et al. 2010; Garbacz et al. 2015; Grabowska et al. 2014) or depletion of nucleotide pools by hydroxyurea (Northam et al. 2010). In this scenario, the recruitment of TLS machinery requires ubiquitylation of PCNA (Northam et al. 2006). In most cases, the TLS-driven spontaneous mutations presumably happen during S-phase replication, because MMR deficiency or defective proofreading elevates mutation rates in strains with impaired replicative pols or accessory factors (Pavlov et al. 2001b; Grabowska et al. 2014; Kiktev et al. 2021).

We demonstrated that elevated production of the pol ζ/Rev1 complex, pol ζ_5_, exerts a spontaneous mutator effect independent of PCNA modification, but is drastically elevated with the addition of hydroxyurea or Cd^2+^. HU depletes dNTP pools and thus suppresses replication. Unlike replicative pols, pol ζ_5_ tolerates low pools (Kochenova et al. 2017), so its participation would alleviate replication inhibition. Elevated mutation rates with HU under conditions of elevated pol ζ_5_ are consistent with this mechanism, although the participation is not benign, as we detected a decline in cell viability. Cd^2+^ partially inhibits MMR; therefore, the synergy suggests that pol ζ_5_-induced mutations arise during replication, as MMR competes with primer extension (Klassen et al. 2023). The combination of elevated pol ζ_5_ and a complete MMR defect in *mlh1-Δ* strains leads to further elevation of mutation rates. We did not see the exact multiplicity as expected from pathways working in series (Morrison et al. 1993). This is because some classes of mispairs (e.g., C-C mispairs) are poor substrates for MMR.

Based on the above observations, we conclude that inaccurate pol ζ_5_, which is naturally devoid of proofreading, directly competes with replicative pols and takes over DNA synthesis when replicative pols dissociate, “stutter”, which can happen, for example, during processing of Okazaki fragments, collision with RNA pols, and in termination zones. It is well documented that pol ζ-dependent mutations induced when normal replication complexes are impaired also occur during S-phase because they pol proofreading corrects them (Garbacz et al. 2015; Grabowska et al. 2014). Future analysis of genomic distribution of mutations by pol ζ_5_ will reveal which regions/motifs are vulnerable.

### Mechanisms of GC pair mutagenesis

The catalytic activity of the Rev3 subunit of pol ζ_5_ and the co-expression of Rev1 were absolutely required for the elevated spontaneous mutagenesis. However, loss of the catalytic transferase activity of Rev1 (Nelson et al. 1996) caused only a partial reduction in overall mutagenesis but a severe reduction in GC→CG transversions, suggesting that Rev3 polymerase activity is required for most mutagenesis, whereas Rev1 catalytic activity contributes predominantly to this specific mutation class.

The profound generation of GC→CG transversions by Rev1 is particularly intriguing because both yeast Rev1 and human REV1 misinsert dCMP opposite template C with efficiencies similar to, or even lower than, those observed opposite template A or T, and approximately 100-fold lower than opposite template G (Haracska et al. 2002; Zhang et al. 2002; Stolyarenko et al. 2024). Pol ζ alone creates CG to GC transversions by making both C:dCMP and G:dGMP mispairs, but these mutations are not a predominant class of its errors (Zhong et al. 2006; Kochenova et al. 2017; Chen and Sugiyama 2017). It is likely that the dominance of CG to GC in mutation spectra in strains with elevated pol ζ_5_ is shaped by MMR, because C:dCMP misincorporation leading to C: C mismatch is one of its less favorable substrates (Brown and Jiricny 1988; Kramer et al. 1989; Lühr et al. 1998).

We detected about two times more C to G than G to C transversions in the coding strand of the *CAN1* gene (despite there being slightly more Gs in the strand (21.1% Gs and 20.0% Cs, **sFig. 5**). The left distal part of chromosome V possesses three replication origins of unknown efficiency to the left of *CAN1*, ARS502, ARS503, ARS504, and ARS504.2; the latter is 12.6 kb from *CAN1.* The closest origin to the left is an early, strong ARS507, 22.6 kb to the right (**sFig**. **9**). The physical mapping of replication suggests that the *CAN1* region is predominantly replicated by replication forks moving towards chromosome end from the centromere because of the activity of ARS507 (McGuffee et al. 2013) (**sFig. 9**). Because most mutations that we see result from dCTP transferase activity of Rev1 creating C-dCMP mispairs we can estimate that the is a bias for the activity of pol ζ/Rev1 complex during lagging DNA strand replication. To firmly confirm this assignment, additional mapping of replication in our strain and under these conditions is necessary, though the bias for the lagging strand is easier to explain.

It is of interest that pol ζ-dependent spontaneous mutagenesis could be PCNA-ubiquitylation dependent (Northam et al. 2006) or independent (this work); however, hotspots of mutations partially overlap, as we found that 50% of them in the *pol3-Y708* strain coincided with mutation hotspots with elevated pol ζ, e.g., at positions 691, 1175, 1195, 1214 in the *CAN1* gene. Shared sites indicate a similar mechanism of pol ζ_5_ operation at certain DNA sequence contexts.

The strongest hotspots for pol ζ-induced GC→CG transversions, defined as sites harboring more than 10 mutations in the combined pol ζ and pol ζ₅Rev3ΔC spectra, shared a common sequence context. The mutated C was usually flanked by A or T at −1, A at +1, and A or T at +2. This pattern suggests that Rev1 preferentially generates GC→CG transversions in AT-rich DNA and may be influenced by the identity of the preceding incorporated nucleotide. Consistent with this notion, analysis of all mutation sites showed that dCMP misincorporation occurred predominantly at template positions containing at least two neighboring purines (Fig. 6C). Further experiments will be necessary to explain the mechanistic basis of this sequence preference.

### Amino acid changes differentially affect pol ζ_5_ and its Rev3CTD variant (***Fig 8***)

Previously, we purified pol ζ variants as described (Fortune et al. 2005; Makarova et al. 2012; Siebler et al. 2022) and found that the preparation of pol ζ_4_ shows four subunits, and the preparation of pol ζ_5_Rev3_ΔC_ shows only two bands, Rev3 and Rev7(Siebler et al. 2022). Pol ζ_5_Rev3_ΔC_ does not co-purify with Pol31/Pol32 under standard conditions. Both pols could extend the primer similarly in primer extension experiments, but primer extension required much more pol ζRev3_ΔC_. On the template with the abasic site, ζRev3_ΔC_ was much less potent than pol ζ_4_, though it inserted opposite an abasic site and extended afterward, allowing it to perform characteristic pol ζ transactions. Low activity pol ζRev3_ΔC_ *in vitro* contrasts with the fact that strains with Rev3-ΔC are proficient in induced mutagenesis (Siebler et al. 2014; Siebler et al. 2022). Strains with inactive CTD lacking Fe-S binding residues are immutable, and pol ζ purifies as a two-subunit complex (Baranovskiy et al. 2012; Kochenova et al. 2017; Makarova et al. 2014). Either the disorganization of CTD by mutations affecting Fe-S cluster binding (**Fig. 2A**) or the complete absence of CTD may make the holoenzyme less stable, and it disassembles during purification. In our work we examined the limit of stability of this structure by introducing amino acid changes affection the interaction with Rev7 dimer, the core component responsible for the assembly of pol ζ_5_ (Malik et al. 2024; Malik et al. 2020; Arianna and Korzhnev 2024). We found that elevated levels of pol ζ/Rev1 can mitigate defects in the wild-type complex but cannot rescue complexes carrying a Rev3 CTD truncation

**Fig. 8.**
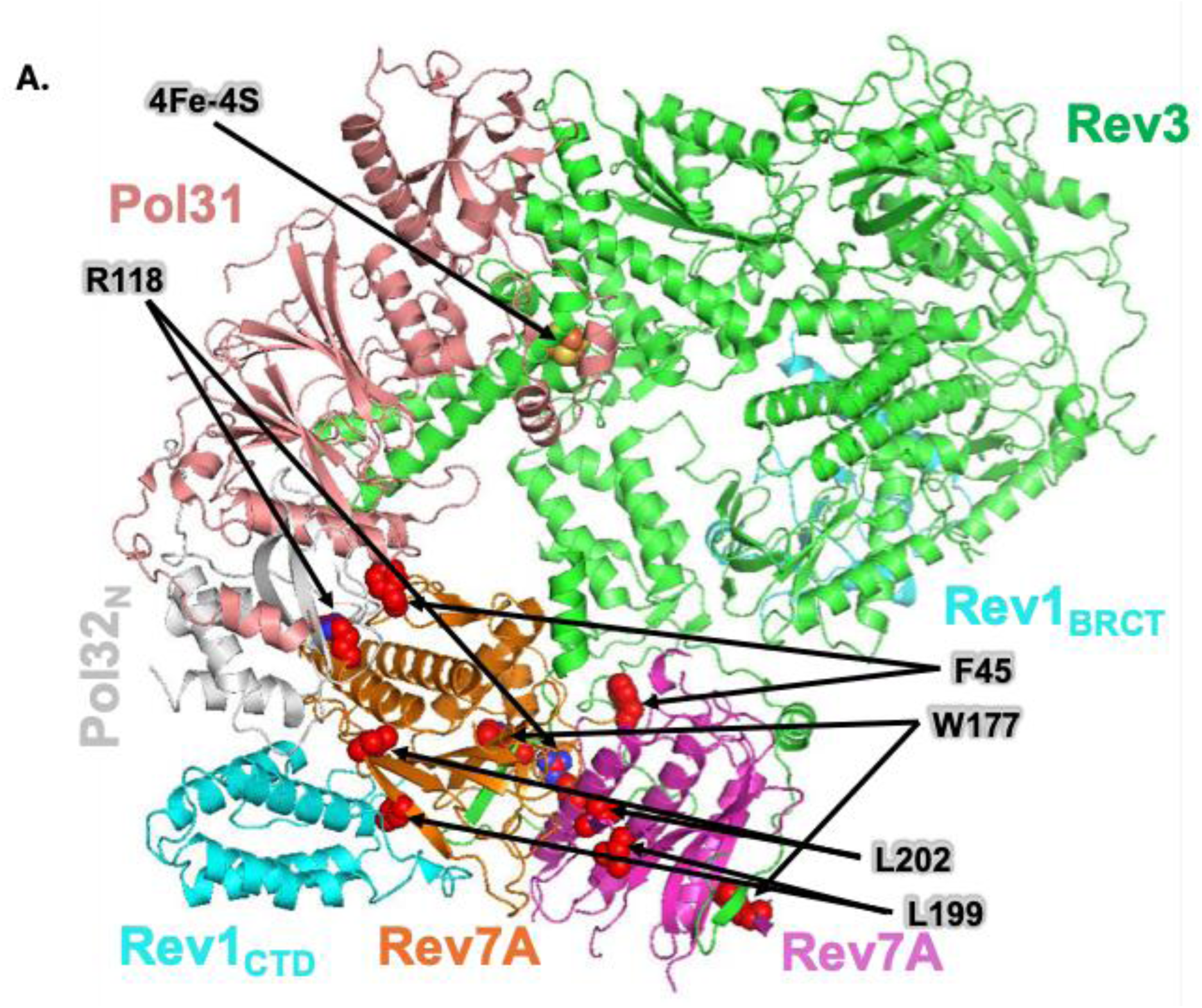
Structural information on pol ζ variants studied. The positions of amino acid substitutions introduced into Rev7 are mapped onto the pol ζ₅ structure using the PDB models **8TLQ** and **8TLT**. Subunits are colored consistently and labeled accordingly. The Fe-S cluster is indicated by an arrow. Residues targeted for mutagenesis within the Rev7 dimer are shown as spheres and identified by arrows extending from their corresponding labels.

Our first step was the construction of strains expressing Rev7-W177A (a change equivalent to W171 in the human homolog of Rev7, **Fig. 8)**. In the crystal structure of the human REV3L fragment and REV7, W171 is one of the critical residues responsible for interaction with partners (Hara et al. 2010). In the cryo-EM structure of yeast pol ζ, homologous W177S in Rev7_A_ and _B_ subunits interact with Rev3 (Malik et al. 2020; Malik et al. 2024). The *rev7-W177A* mutation moderately lowered induced mutagenesis under standard conditions. However, under overexpression, the mutagenesis reached wild-type level (**Table 2A, first row**). Strikingly, the same allele completely abolished induced and spontaneous mutagenesis in yζ_5_Rev3_ΔC_ strains (**Table 2B, first row)**. We propose that, in the presence of the Rev3 CTD, the W177A change does not severely affect pol ζ complex assembly because the dominant interaction is with the Rev3 CTD. When the CTD is missing, assembly occurs via weaker interactions, and a single W177A change is sufficient to destabilize the complex altogether, regardless of expression level.

Most amino acid changes (except L202A alone) affecting interfaces of Rev7 interacting with Pol32 and Rev1 had similar effects, leading to mild or no effects when present in pol ζ_5_ (**Table 2A**) but led to a complete ablation of spontaneous mutagenesis by elevated pol ζ (**Table 2B**). ΘΩΡ-ιnduced mutagenesis effects were seen under low pol ζ_5_ expression but disappeared at high expression. The strains with changes in yζ_5_Rev3_ΔC_ were UV-immutable at low expression (except L202A, which showed a substantial reduction), but were much more sensitive at high expression. Interestingly, F45 and the double F45A L202A retained 25-50% of UV mutagenesis despite abolishing spontaneous mutagenesis, suggesting a difference in the mechanism of mutation appearance. Further work is required to determine whether this interaction is most critical for the function of yζ_5_Rev3_ΔC_ in vivo. We conclude that yζ_5_Rev3_ΔC_ is a minimal mutasome with adverse vulnerability to any further protein sequence changes.

### Mutational signatures in cancer

The prevalent mutations induced by Pol ζ were C•G to C•G transversions. The pol ζ/Rev1 mutagenesis signature coincides with several CG-to-GC COSMIC signatures of unknown etiology in cancer. We propose that these signatures arise from elevated TLS in tumors. Overexpression of specialized pols characterizes many cancers (Albertella et al. 2005) and is considered the source of mutations that cause some cancers (Bainbridge and Daigaku 2024). In addition, TLS polymerases induce collateral mutations after therapy that contribute to tumor mutation landscapes (Póti et al. 2022).

Several cancer mutation signatures (Alexandrov et al. 2020; Alexandrov et al. 2013a; Alexandrov et al. 2013b) including SBS13, SBS39, SBS86, and SBS103 display a similar predominance of G to C transversion. However, a cosine similarity score exceeding 0.7 was observed only for SBS39 (**Fig. 7, sTable 6**). This relatively rare signature of unknown etiology appears to represent a late-acting mutational process, being enriched in late clonal mutations, suggesting increased activity of signature-causing factors during tumor evolution (Torrens et al. 2025). The signature was detected various types of cancer, e.g., head and neck, pancreatic, central nervous system medulloblastoma, ovarian, and breast cancer, especially with BRCA1/2 defects (Ferrer-Torres et al. 2025), however, it does not correlate well with the *bona fide* homologous recombination defect signature, SBS3. Thus, is unclear whether this correlates with the known role of pol ζ in recombination (Introduction). Based on our finding that Pol ζ induces mutations by competing with replicative polymerases, we propose that replication-stress conditions in cancer cells promote Pol ζ involvement in S-phase replication, forming this characteristic signature. Indeed, SBS39 correlates better with mutations found in early than in late replication regions in the COSMIC database, though it shows no leading/lagging strand bias (Tate et al. 2019).

SBS86 and SBS103 are poorly understood mutational signatures with limited characterization in cancer genomes. According to the COSMIC signature catalogue, SBS103 is currently classified as having unknown etiology and lacks clear biological associations in the literature.

SBS86 has been detected at low prevalence across multiple tumor types, including hematologic cancers, but its origins remain unclear. It appears in a subset of relapsed acute lymphoblastic leukemia (ALL) cases, reportedly affecting around 10% of relapsed ALL patients and enriched in certain subtypes such as hyper-diploid ALL (van der Ham et al. 2024), and is noted along with other treatment-associated signatures in therapy-exposed cohorts. However, definitive causal processes have not yet been established (Van der Ham et al. 2025). Weak resemblance to our spectra may reflect general properties of pol ζ-associated DNA transactions.

One paradoxical observation is the weak resemblance between the mutasome spectra in yeast and SBS13, a mutational signature attributed to the activity of APOBEC cytosine deaminases. Because yeast lacks these DNA-editing enzymes, this connection initially appears counterintuitive. However, this CG-to-GC transversion signature is generated by REV1 activity during the processing of deaminated uracil (Taylor et al. 2013; Harris 2015); therefore, the context is determined by the overlap of APOBEC, base excision repair, and REV1 specificities. Apparently, trinucleotide mutational specificity analysis captured the Rev1 component of SBS13, albeit with lower confidence than SBS39.

## Acknowledgement

We thank Elizabeth Moore for excellent technical assistance and Aileen Martin for help preparing mutation spectra figures. The work was funded by in part by NE DHHS grant 2023-46. Research part done at UNMC was also supported by the National Cancer Institute under award number P30 CA036727. The content is solely the responsibility of the authors and does not necessarily represent the official views of the National Cancer Institute.

