## Supplemental materials for "Minimal eukaryotic DNA polymerase ζ/Rev1 GC-pair mutasome"

Supplemental Table 1. Main oligonucleotide primers used in the study*.

| name | sequence 5' to 3'¶ | application |
|---|---|---|
| gal-11,571R | CTCCGAAGGAAGACTCT | Verification of integration of REV3rc into chromosome, gene start |
| rev3rc-678R | TATGGCAAAATACCGTGA | |
| rev3rc-4501F | ATTGGACAGATTGTTCAA | Verification of integration of *REV3rc* or *rev3rc*-Δ into chromosome, gene end |
| rev3rc-5178R | GAATTGGAGCTCCACCGC | |
| rev3rc-NTN-F | GTTTACGGT**AAC**ACT**AAC**TCTTTGTTCGTTTAC | SDM to create **catalytic dead** Rev3 |
| rev3rc-NTN-R | CAAAGA**GTT**AGT**GTT**ACCGTAAACAACCTTAGC | |
| rev3rc-cat seq | GGACGTTAGAGTTATGAT | Sequencing of the region encompassing *REV3* catalytic site |
| gal-11,571R | CTCCGACGGAAGACTCTC | Verification of integration of *REV1rc* into chromosome, gene start |
| rev1rc-7083F | AGAGATGTTCTTAGTGGTGA | |
| rev1rc-4036F | CGGGCCTCTTCGCTATTA | Verification of integration of *REV1rc* into chromosome, gene end |
| rev1rc-4635R | CGTTAAGTTGTTTGTGAAGT | |
| R1 M1-Del F | AGCTTCTCTAGTGTAAACGGTGTTC | |

| | | |
|---|---|---|
| R1 M1-Del R | AAGGATCAGTACAAGGACCAGATC | SDM to create M1 mutant of Rev1 |
| Rev1 6700F | ATTCTCTTCTTCAATGGCAA | Verification of sequence of M1 mutant |
| rev1rc-de->aa-F | ACGCAAACAGC**TGCCGC**AATGGAGATTGGC | SDM to create allele encoding for the **catalytic dead** Rev1 |
| rev1rc-de->aa-R | CCATT**GCGGCA**GCTGTTTGCGTTAGAATCATC | |
| rev1rc-5743F | GACAAGTTGGACTTAAAGGT | Sequencing of the region encompassing *REV1* catalytic site |
| gal-to rev7-F | GGTGGTAATGCCATG | Verification of integration of *REV7rc* into chromosome, gene start |
| rev7rc-302R | AGATTGGTAAGTAGTGTA | |
| rev7rc-Cend-F | GTGACGACAAGATTTTGAACGG | Verification of integration of *REV7rc* into chromosome, gene send |
| rev7rc-1202-R | CAGCTATGACCATGATTA | |
| rev7rc-W177A-F | GAGACTCTAAC**GCG**GTTAAGTGTCAAGAAGACG | SDM to create Rev7 encoding for the W177A change |
| rev7rc-W177A-R | GACACTTAAC**CGC**GTTAGAGTCTCTTTCAATTTCAG | |
| rev7rc-451-F | TTGAAAAGTACGTTTTGG | Sequencing of the C-terminal half of *REV7* |
| DIS-REV3F-nat | TATTTGAGTCAATACAAAACTACAAGTTGTGGCGAAATAAAATGTTTGGAA**CGTACGCTGCAGGTCGAC** # | Chromosomal *REV3* deletion |

| | | |
|---|---|---|
| | | by substitution of ORF by *nat* module (nourseothricin resistance) |
| DIS-REV3R-nat | TAGAAACAAATAACTACTCATCATTTTGCGAGACATATCTGTGTCTAGATTA**TCGATGAATTCGAGCTCG** # | |
| REV3 left-0 | ACCATTGTCCAAAGCTGTCG | Verification of deletion of *REV3* |
| REV3 right-0 | AGGTTCAAACGGAGGAGGAA | |
| REV1dis::ade2F | GATTTTCTCAAAATAAATCGATACTGCATTTCTAGGCATATCCAGCG**ATGAACTGTATCGAAACGTT**# | Chromosomal *REV1* deletion by substitution of ORF by the *ADE2* module (selection for Ade$^+$) |
| REV1dis::ade2R | CAAACTGCGTGTTTACTGTATGCTGAAATGTTTTTTTTTTTTTAA**GGAAATTCCAATTATCCCGA**# | |
| REV1ver633F | TCGCAAATTACTCATCACAT | Verification of deletion of *REV1* |
| REV1ver4899R | AGCAAGGGTTGGATAATTTT | |
| REV7_155F | AAGCCTCTCTTTATTTCGTC | Verification of *REV7* deletion |
| REV7_1689R | AACACCATCACCATATAACG | |

*Sequences of primers use for sequencing of the pol ζ-encoding genes and other not listed in the Table are available upon request.

¶ In case of SDM, codons encoding for a new amino acid are in **bold**.

### in case of primers for one-step gene disruption the part necessary for amplification of marker gene used for selection is in **bold**.

Supplemental Table 2. Principal yeast strain used in the study*.

| Catalog name** | Other name | genotype | reference |
|---|---|---|---|
| yo1636 | yJF1 | *MAT***a** *ura3-1 trp1-1 his3-11,15 ade2-1 can1-100 leu2-3,112 bar1::hph pep4Δ::kanMX rad5-G535R* | {Yeeles, 2015 #5} |
| yo1155 | α LAN201-Δura3 | *MATα ura3-Δ trp1-289 his7-2 ade5-1 CAN1 leu2-3,112 lys2-Tn5-13* | {Stepchenkova, 2017 #51} |
| yo1318 | YPOM600 | Diploid resulting from mating yJF1 to α LAN201-Δura3 | This study |
| yo1979 /yo2990 | 1C-YPOM616 R5$^{+}$ | *MAT***a** *ura3-1 trp1-1 his3-11,15 ade2-1 CAN1 leu2-3,112 bar1Δ::hph pep4Δ::kanMX RAD5* | This study |
| yo1982 /yo2992 | 1C-YPOM616 R5$^{+}$ rev3Δ | *MAT***a** *ura3-1 trp1-1 his3-11,15 ade2-1 CAN1 leu2-3,112 bar1Δ::hph pep4Δ::kanMX RAD5 rev3Δ::nat* | This study |
| yo2993 | 1C-YPOM616 R5$^{+}$ rev7Δ | *MAT***a** *ura3-1 trp1-1 his3-11,15 ade2-1 CAN1 leu2-3,112 bar1Δ::hph pep4Δ::kanMX RAD5 rev7Δ* | This study |
| yo2815 | 1C-YPOM616 R5$^{+}$ *rev1Δ:A* | *MAT***a** *ura3-1 trp1-1 his3-11,15 ade2-1 CAN1 leu2-3,112 bar1Δ::hph pep4Δ::kanMX RAD5 rev1Δ::ADE2* | |
| yo3017 | 1C-YPOM616 R5$^{+}$ rev37Δ | *MAT***a** *ura3-1 trp1-1 his3-11,15 ade2-1 CAN1 leu2-3,112 bar1Δ::hph pep4Δ::kanMX RAD5 rev3Δ::nat rev7Δ* | This study |
| yo3070 /yo3091 | 1C-YPOM616 R5$^{+}$ Δr371 | *MAT***a** *ura3-1 trp1-1 his3-11,15 ade2-1 CAN1 leu2-3,112 bar1Δ::hph pep4Δ::kanMX RAD5 rev3Δ::nat rev7Δ rev1Δ::ADE2* | This study |
| yo2631 | $y\zeta_{5*v1}$ | *MAT***a** *ura3-1_REV3rc_POL31rc_URA3 trp1-1_REV1rc_TRP1 his3-11,15_REV7rc_POL32rc_CBP_HIS4 ade2-1 CAN1 leu2-3,112 bar1Δ::hph pep4Δ::kanMX RAD5 rev3Δ::nat REV1 REV7* | This study |
| yo2635 | $y\zeta_{5}Rev3_{\Delta Cv1}$ | *MAT***a** *ura3-1_REV3rc_ΔC_6xHis_POL31rc_URA3 trp1-1_REV1rc_TRP1 his3-11,15_REV7rc_POL32rcCBP_HIS4 ade2-1 CAN1 leu2-3,112 bar1Δ::hph pep4Δ::kanMX RAD5 rev3Δ::nat REV1 REV7* | This study |

| | | | |
|---|---|---|---|
| yo2703 | y$\zeta_{5*_PCNAK164R}$ | *MAT***a** *ura3-1_REV3rc_POL31rc_URA3 trp1-1_REV1rc_TRP1 his3-11,15_REV7rc_POL32rc_CBP_HIS4 ade2-1 CAN1 leu2-3,112 bar1Δ::hph pep4Δ::kanMX RAD5 rev3Δ::nat REV1 REV7 pcnaK164A-LEU2* | This study |
| yo2700 | y$\zeta_5$Rev3$_{\Delta C_PCNAK164R}$ | *MAT***a** *ura3-1_REV3rc_ΔC_6xHis_POL31rc_URA3 trp1-1_REV1rc_TRP1 his3-11,15_REV7rc_POL32rcCBP_HIS4 ade2-1 CAN1 leu2-3,112 bar1Δ::hph pep4Δ::kanMX RAD5 rev3Δ::nat REV1 REV7pcnaK164A-LEU2* | |
| yo3111 | y$\zeta_{5v2}$ | *MAT***a** *ura3-1_REV3rc_POL31rc_URA3 trp1-1_REV1rc_TRP1 his3-11,15_REV7rc_POL32rc_CBP_HIS4 ade2-1 CAN1 leu2-3,112 bar1Δ::hph pep4Δ::kanMX RAD5 rev3Δ::nat rev1Δ::ADE2 rev7Δ* | |
| yo3118 | y$\zeta_5$Rev3$_{\Delta Cv2}$ | *MAT***a** *ura3-1_REV3rc_ΔC_6xHis_POL31rc_URA3 trp1-1_REV1rc_TRP1 his3-11,15_REV7rc_POL32rcCBP_HIS4 ade2-1 CAN1 leu2-3,112 bar1Δ::hph pep4Δ::kanMX RAD5 rev3Δ::nat rev1Δ::ADE2 rev7Δ* | This study |
| yo3238 | E134 | *MATα ura3-52 trp1-289 his7-2 ade5-1 CAN1 leu2-3,112 lys2::InsE*$_{A14}$ | {Tran, 1999 #229;Shcherbakova, 1999 #175} |
| yo3238 | E134 *pol2-4* | same, but *pol2-4* | Constructed in this study as described in {Morrison, 1991 #130} |
| yo3239 | E134 *pol2-V411L* | same, but *pol2-V411L* | Obtained from Dr. Shcherbakova, described in {Barbari, 2022 #226} |
| yo3242 | E134 *pol3-5DV* | same, but *pol3-5DV* | Constructed in this study as described in {Jin, 2005 #230} |

+ The Table shows only main strains; during the work we constructed strains with various assortment of natural and recoded genes and alleles encoding pol ζ/Rev1 (pol $\zeta_5$) components and their variants to examine the effects of mutations in individual genes. Their genotypes and names are available upon request.

** Entries in Pavlov laboratory strain collection. Two numbers are given with slash mean same strain stored at different time, larger number denotes a more recent clone stored.

sTable 3A. Effects of expression of pol ζ components

| Wild-type Rev3 catalytic subunit in pol $\zeta_5$ | | | | | | | |
|---|---|---|---|---|---|---|---|
| Integrated pol ζ component gene | Parent strain | Mutagenesis in comparison to wild-type strain | | | | | |
| | | Low expression | | | High expression | | |
| | | Spontaneous mutator effect | Complementation of UVR mutagenesis defect | | Spontaneous mutator effect | Complementation of UVR mutagenesis defect | |
| | | | Low dose | High dose | | Low dose | High dose |
| none | *rev3-Δ* | 1/2 | 0 | 0 | 1/2 | 0 | 0 |
| *REV3rc POL31rc* | *rev3-Δ* | 1 | 1 | 1 | 1 | 1 | 1 |
| *REV7rc POL32rc* | *rev7-Δ* | 1 | 1 | 1 | 1 | 1 | 1 |
| *REV1rc* | *rev1-Δ* | 1 | 1 | 1 | 1 | 1 | 1 |
| *REV3rc POL31rc REV7rc POL32rc* | *rev3-Δ rev7-Δ* | 1 | 1 | 1 | 1 | 1 | 1 |
| *REV3rc POL31rc REV7rc POL32rc REV1rc* | *rev3-Δ rev7-Δ rev1-Δ* | 1 | 1 | 1 | 20 | 1 | 1 |
| Same but *rev3_rev3-C*1446, 1449A, 1468A 1473A | *rev3-Δ rev7-Δ rev1-Δ* | 1/2 | 0 | 0 | 20 | 1 | 1 |

sTable 3B. Effects of expression of pol ζ components.

| CTD-less Rev3 catalytic subunit in pol $\zeta_5 Rev3_{\Delta CTD}$ | | | | | | | |
|---|---|---|---|---|---|---|---|
| Integrated pol ζ component gene | Parent strain | Mutagenesis in comparison to wild-type strain | | | | | |
| | | Low expression, fold change in mutagenesis | | | High expression, fold change in mutagenesis | | |
| | | Spontaneous mutator effect | Complementation of UVR mutagenesis defect | | Spontaneous mutator effect | Complementation of UVR mutagenesis defect | |
| | | | Low dose | High dose | | Low dose | High dose |
| none | *rev3-Δ* | 1/2 | 0 | 0 | 1/2 | 0 | 0 |
| *REV3rc-ΔC POL31rc* | *rev3-Δ* | 1 | 1 | 1/2 | 1 | 1 | 1 |
| *REV7rc POL32rc* | *rev7-Δ* | 1 | 1 | 1 | 1 | 1 | 1 |
| *REV1rc* | *rev1-Δ* | 1 | 1 | 1 | 1 | 1 | 1 |
| *REV3rc-ΔC POL31rc REV7rc POL32rc* | *rev3-Δ rev7-Δ* | 1 | 1 | 1/2 | 1 | 1 | 1 |
| *REV3rc-ΔC POL31rc REV7rc POL32rc REV1rc* | *rev3-Δ rev7-Δ rev1-Δ* | 1 | 1 | 3/4 | 20 | 1 | 1 |

Supplementary Table 4. Summary of spontaneous mutation rates and UVR-induced mutant frequencies in yeast strains with pol ζ variants.

| Critical change | Function / interaction affected | Without induction | UVR, J/m$^2$ ± SE | | Induced | UVR, J/m$^2$ | |
|---|---|---|---|---|---|---|---|
| | | Spontaneous* | 25** | 50** | Spontaneous* | 25** | 50** |
| **Changes in yζ$_5$** | | | | | | | |
| - | | 2.6 | 78 | 193 | 20.1 | 61 | 167 |
| | | 1.4-3.4 | ± 25 | ± 44 | 17.1-25.3 | ± 22 | ± 51 |
| *rev3$_{cat}$* | null Rev3 activity | 1.2 | 2.0 | n/a[#] | 2.9 | 2.1 | 1.3 |
| | | 0.8-1.7 | ± 0.9 | ± 0.8 | 1.7-3.5 | ± 0.5 | ± 1.1 |
| *rev3-C*1446, 1449A, 1468A 1473A | Inability of Rev3's CTD to bind Fe-S cluster | 1.1 | 2.3 | n/a[#] | 28.7 | 80 | 143 |
| | | 0.5-1.4 | ± 1.9 | | 25.0-46.0 | ± 29 | ± 63 |
| *rev1$_{cat}$* | null Rev1's transferase activity | 2.7 | 60 | 178 | 6.9 | 71 | 180 |
| | | 1.5-4.4 | ± 21 | ± 51 | 1.1-8.9 | ± 15 | ± 66 |
| *rev1-ΔM1* | Elevated activity of Rev1 *in vitro* | 2.5 | 67 | 179 | 19.9 | 79 | 172 |
| | | 1.6-3.7 | ± 25 | ± 34 | 14.2-24.9 | ± 32 | ± 49 |
| *rev7-W177A* | Rev3 and Rev1 | 2.1 | 12 | 11 | 19.2 | 87 | 230 |
| | | 1.4-4.1 | ± 7 | ± 6 | 9.1-15.6 | ± 24 | ± 84 |
| *rev7-L199A* | CTD of Rev1 | 2.5 | 35 | 70 | 18.4 | 93 | 188 |
| | | 1.7-3.2 | ± 11 | ± 24 | 17.2-33.3 | ± 29 | ± 41 |
| *rev7-L202A* | CTD of Rev1 | 2.7 | 69 | 165 | 24.3 | 102 | 177 |
| | | 0.7-5.2 | ± 9 | ± 44 | 18.9-28.4 | ± 39 | ± 46 |
| *rev7-F45A* | NTD of Pol32 | 1.4 | 85 | 204 | 15.2 | 99 | 246 |
| | | 0.5-1.6 | ± 17 | ± 77 | 12.2-22.9 | ± 18 | ± 87 |
| *rev7-R118A* | NTD of Pol32 | 1.5 | 53 | 83 | 23.2 | 59 | 136 |
| | | 0.8-3.1 | ± 15 | ± 19 | 18.1-33.4 | ± 8 | ± 39 |

| | | | | | | | |
|---|---|---|---|---|---|---|---|
| *rev7-F45A L202A* | NTD of Pol32 and CTD of Rev1 | 1.7 | 26 | 115 | 12.2 | 59 | 152 |
| | | 0.9-2.9 | ± 8 | ± 25 | 9.3-15.6 | ± 32 | ± 48 |
| **Changes in yζ$_5$Rev3$_{\Delta C}$** | | | | | | | |
| - | | 2.0 | 61 | 3.6 | 21.0 | 54 | 159 |
| | | 0.7-4.7 | ± 12 | ± 4 | 13.0-34.1 | ± 15 | ± 46 |
| *rev3$_{cat}$* | null Rev3 activity | 1.6 | 3.1 | 1.9 | 3.1 | 1.8 | 2.9 |
| | | 0.9-1.9 | ± 1.1 | ± 1.5 | 2.1-3.6 | ± 0.7 | ± 1.8 |
| *rev1$_{cat}$* | null Rev1 activity | 1.5 | 45 | 11 | 5.7 | 55 | 157 |
| | | 1.2-3.9 | ± 12 | ± 6 | 4.5-8.2 | ± 12 | ± 52 |
| *rev1-ΔM1* | Elevated activity of Rev1 *in vitro* | 2.3 | 59 | 4.1 | 18.3 | 49 | 166 |
| | | 1.4-2.4 | ± 16 | ± 2.9 | 8.3-25.5 | ± 11 | ± 51 |
| *rev7-W177A* | Rev3 and Rev1 | 1.4 | 1.3 | 4.2 | 1.6 | 1.9 | 2.2 |
| | | 0.9-3.5 | ± 1.2 | ± 2.2 | 1.2-4.2 | ± 0.9 | ± 1.7 |
| *rev7-L199A* | CTD of Rev1 | 2.6 | 3.9 | 4.5 | 2.8 | 4.5 | 2.8 |
| | | 0.9-3.5 | ± 1.3 | ± 1.2 | 0.9-4.2 | ± 2.5 | ± 1.9 |
| *rev7-L202A* | CTD of Rev1 | 2.9 | 35 | 11.7 | 25.2 | 66 | 230 |
| | | 0.8-3.1 | ± 14 | ± 6.2 | 17.2-34.1 | ± 11 | ± 69 |
| *rev7-F45A* | NTD of Pol32 | 0.8 | 5.4 | 2.1 | 2.9 | 29 | 117 |
| | | 0.2-1.6 | ± 1.8 | ± 0.4 | 1.7-3.4 | ± 9 | ± 23 |
| *rev7-R118A* | NTD of Pol32 | 3.2 | 3.2 | 2.3 | 2.2 | 3.3 | 4.1 |
| | | 1.3-3.1 | ± 2.1 | ± 0.8 | 1.5-3.6 | ± 1.7 | ± 2.8 |
| *rev7-F45A L202A* | NTD of Pol32 and CTD of Rev1 | 2.1 | 3.4 | 2.5 | 3.5 | 17 | 11 |
| | | 1.4-2.1 | ± 1.1 | ± 1.9 | 1.4-4.7 | ± 7 | ± 8 |

n/a[#] - precise measurement is difficult because of low viability (<1%) at the dose

Supplemental Table 5. Mutations in the *CAN1* gene induced by expression of pol ζ variants.

**A.** All Mutation types.

| Strain / Mutation Type | $y\zeta_5$ | | $y\zeta_5Rev3_{\Delta C}$ | | $y\zeta_5Rev1_{cat}$ | | $y\zeta_5Rev3_{\Delta C}Rev1_{cat}$ | |
|---|---|---|---|---|---|---|---|---|
| | n | % | n | % | n | % | n | % |
| Transitions | | | | | | | | |
| GC -> AT | 24 | 8.3 | 33 | 9.5 | 35 | 32.7 | 68 | 32.4 |
| AT -> GC | 15 | 5.2 | 21 | 6.0 | 2 | 1.9 | 18 | 8.6 |
| Transversions | | | | | | | | |
| GC -> TA | 40 | 13.8 | 47 | 13.5 | 15 | 14.0 | 71 | 33.8 |
| GC -> CG | 152 | 52.4 | 176 | 50.4 | 29 | 27.1 | 16 | 7.6 |
| AT -> CG | 28 | 9.7 | 31 | 8.9 | 1 | 0.9 | 2 | 1.0 |
| AT -> TA | 18 | 6.2 | 8 | 2.4 | 20 | 18.7 | 22 | 10.5 |
| +/-1 or 2 Indels: | | | | | | | | |
| insertions | 3 | 1.0 | 1 | 0.3 | 2 | 1.9 | 1 | 1.0 |
| deletions | 10 | 3.4 | 31 | 8.9 | 3 | 2.8 | 10 | 4.8 |
| Other* | | | 1 | 0.3 | | | 1 | 0.5 |
| Total mutations | 290 | | 349 | | 107 | | 210 | |
| P** | 0.0284 | | | | | | | |
| | 0.0001 | | | | | | | |
| | 0.0001 | | | | | | | |
| | | | 0.0001 | | | | | |
| | | | 0.0001 | | | | | |
| | | | | | 0.0001 | | | |

*- complex mutations involving several nucleotides or large deletions

**- Fisher exact test

**B.** Spectra comparisons:

| Strain / Mutation Type | wild-type | | $y\zeta_5$ | | $y\zeta_5Rev3_{\Delta C}$ | | $y\zeta_5Rev1_{cat}$ | | $y\zeta_5Rev3_{\Delta C}$ $Rev1_{cat}$ | |
|---|---|---|---|---|---|---|---|---|---|---|
| | n | f* | n | f | n | f | n | f | n | f |
| GC -> CG | 62 | 0.12 | 152 | 0.55 | 176 | 0.56 | 29 | 0.29 | 16 | 0.09 |
| Total single base substitution mutations | 537 | | 277 | | 316 | | 102 | | 197 | |
| P** | $<10^{-40}$ | | | | | | | | | |
| | $<10^{-39}$ | | | | | | | | | |
| | | | $<10^{-10}$ | | | | | | | |
| | | | | | $<10^{-24}$ | | | | | |

* f – fraction

** - chi square test

Supplemental Table 6. Mutation signature of elevated active pol ζ mirrors Cosmic cancer signatures.

| | $y\zeta_5$ | | $y\zeta_5Rev3_{\Delta C}$ | | $y\zeta_5Rev1_{cat}$ | | $y\zeta_5Rev3_{\Delta C}Rev1_{cat}$ | |
|---|---|---|---|---|---|---|---|---|
| **Cosmic signature** | cosine | $P_{cos}$ | cosine | $P_{cos}$ | cosine | $P_{cos}$ | cosine | $P_{cos}$ |
| **SBS39** | 0.73 | 0.001 | 0.55 | 0.0004 | 0.42 | 0.078 | 0.33 | 0.0249 |
| **SBS86** | 0.52 | 0.0029 | 0.37 | 0.0407 | 0.44 | 0.0097 | 0.28 | 0.0951 |
| **SBS103** | 0.59 | 0.0153 | 0.37 | 0.0301 | 0.34 | 0.0482 | 0.31 | 0.886 |
| **SBS13** | 0.45 | 0.0058 | 0.56 | 0.0109 | 0.3 | 0.172 | 0.14 | 0.9155 |

Supplemental Fig 1. DNA sequence information of the constructs dodne in our study.

A. Fragment for *REV7* flanking region for creation chromosomal deletion of the *REV7* gene. We show nucleotide sequences of synthesized fragment with the restriction enzyme recognition sites in the nucleotide sequences of synthesized fragment underlined, Part of fragment corresponding to left part of *REV7* flanking region before ATG codon of ORF is in lowercase letters, right part of fragment after termination codon of ORF is in UPPERCASE.

```
SalI
gtcgacgactaaaattacggaagatacaaacgacaaacgcgctaaaggcaacggaggtgatgattttattatt
gctcacacaactgatgttgctgacagatggcaaatccttttcattttagttattatcattgggtgctttttgg
aaaaaagatgaactaagcctctctttatttcgtccattttattctagcttttacattatgctttcacctttac
caaaaaatagtatgtatttcttttccccttgctataaatcaaaagtacgtagaaacataaacaggcgagaaat
acatatgcatatacgtctacactactaatttaatttccactttcttaatatcagtcaatcgaaaccactagct
ttcacactaagactagaccctgtttaactaatataaacttttccagaaagctaaaaaaaaggtcggaaaaaga
aggtatacgtttttgcgacgactgagatccaagaagaaaaaaaaaaatagtaatcgttgcgtcagcttttTGCAA
AAATGAAATTTGAAGAATGGAATTAAAATTAAATGTCTCTAAGTATAAAAACTAAATAAAATATAGGGTATCT
GTAAACCACTAATAAAAAATAAAACAGGAAAATAAAATTAAGCAAAATATCGTTGCGTATAATCACAGCATTT
CTGGGATGCGTCCGTGAATGTCCTTAACTTGACCAACATTGACATTACTCTTATTTGAAAAATCTACTAACCT
TTTCTTACTTGGTCTTCTGTCTGGTTCCATGCTATGGACCCAGCAAAAATTTTCACCATCTTTAACCGGAACA
AAATCGTCAGAAGATGAAGTTGACATTGTTGTGGGAGTGATTCCGTTTTTACCGCTTTGAGAGTCTTGAATAT
TTTGTAAGTGGCGGTTACGATGCTTCGTTACGTTATATGGTGATGGTGTTACTGATTTTCTTACGGGAGAAGG
GCTAATATTGCTGTTCCAAGGGTCCAGTGAAGACATAGTGACATCCCTTGACGTACTCTAGA
    XbaI
```

B. Recoded gene fragments. Nucleotide sequences of newly synthesized optimized genes encoding components of pol $\zeta_5$.
The restriction enzyme recognition sites are underlined, Sites Open reading frames are in capital letters with ATG and stop codons in **bold**.

*REV3rc*:

SgrAI
caccggtgtataaaaca**ATG**TCTAGAGAATCTAACGACACTATTCAATCTGACACTGTTAGATCTTCTT
CTAAGTCTGACTACTTCAGAATTCAATTGAACAACCAAGACTACTACATGTCTAAGCCAACTTTCTTGG
ACCCATCTCACGGTGAATCTTTGCCATTGAACCAATTCTCTCAAGTTCCAAACATTAGAGTTTTCGGTG
CTTTGCCAACTGGTCACCAAGTTTTGTGTCACGTTCACGGTATTTTGCCATACATGTTCATTAAGTACG
ACGGTCAAATTACTGACACTTCTACTTTGAGACACCAAAGATGTGCTCAAGTTCACAAGACTTTGGAAG
TTAAGATTAGAGCTTCTTTCAAGAGAAAGAAGGACGACAAGCACGACTTGGCTGGTGACAAGTTGGGTA

ACTTGAACTTCGTTGCTGACGTTTCTGTTGTTAAGGGTATTCCATTCTACGGTTACCACGTTGGTTGGA
ACTTGTTCTACAAGATTTCTTTGTTGAACCCATCTTGTTTGTCTAGAATTTCTGAATTGATTAGAGACG
GTAAGATTTTCGGTAAGAAGTTCGAAATTTACGAATCTCACATTCCATACTTGTTGCAATGGACTGCTG
ACTTCAACTTGTTCGGTTGTTCTTGGATTAACGTTGACAGATGTTACTTCAGATCTCCAGTTTTGAACT
CTATTTTGGACATTGACAAGTTGACTATTAACGACGACTTGCAATTGTTGTTGGACAGATTCTGTGACT
TCAAGTGTAACGTTTTGTCTAGAAGAGACTTCCCAAGAGTTGGTAACGGTTTGATTGAAATTGACATTT
TGCCACAATTCATTAAGAACAGAGAAAAGTTGCAACACAGAGACATTCACCACGACTTCTTGGAAAAGT
TGGGTGACATTTCTGACATTCCAGTTAAGCCATACGTTTCTTCTGCTAGAGACATGATTAACGAATTGA
CTATGCAAAGAGAAGAATTGTCTTTGAAGGAATACAAGGAACCACCAGAAACTAAGAGACACGTTTCTG
GTCACCAATGGCAATCTTCTGGTGAATTCGAAGCTTTCTACAAGAAGGCTCAACACAAGACTTCTACTT
TCGACGGTCAAATTCCAAACTTCGAAAACTTCATTGACAAGAACCAAAAGTTCTCTGCTATTAACACTC
CATACGAAGCTTTGCCACAATTGTGGCCAAGATTGCCACAAATTGAAATTAACAACAACTCTATGCAAG
ACAAGAAGAACGACGACCAAGTTAACGCTTCTTTCACTGAATACGAAATTTGTGGTGTTGACAACGAAA
ACGAAGGTGTTAAGGGTTCTAACATTAAGTCTAGATCTTACTCTTGGTTGCCAGAATCTATTGCTTCTC
CAAAGGACTCTACTATTTTGTTGGACCACCAAACTAAGTACCACAACACTATTAACTTCTCTATGGACT
GTGCTATGACTCAAAACATGGCTTCTAAGAGAAAGTTGAGATCTTCTGTTTCTGCTAACAAGACTTCTT
TGTTGTCTAGAAAGAGAAAGAAGGTTATGGCTGCTGGTTTGAGATACGGTAAGAGAGCTTTCGTTTACG
GTGAACCACCATTCGGTTACCAAGACATTTTGAACAAGTTGGAAGACGAAGGTTTCCCAAAGATTGACT
ACAAGGACCCATTCTTCTCTAACCCAGTTGACTTGGAAAACAAGCCATACGCTTACGCTGGTAAGAGAT
TCGAAATTTCTTCTACTCACGTTTCTACTAGAATTCCAGTTCAATTCGGTGGTGAAACTGTTTCTGTTT
ACAACAAGCCAACTTTCGACATGTTCTCTTCTTGGAAGTACGCTTTGAAGCCACCAACTTACGACGCTG
TTCAAAAGTGGTACAACAAGGTTCCATCTATGGGTAACAAGAAGACTGAATCTCAAATTTCTATGCACA
CTCCACACTCTAAGTTCTTGTACAAGTTCGCTTCTGACGTTTCTGGTAAGCAAAAGAGAAAGAAGTCTT
CTGTTCACGACTCTTTGACTCACTTGACTTTGGAAATTCACGCTAACACTAGATCTGACAAGATTCCAG
ACCCAGCTATTGACGAAGTTTCTATGATTATTTGGTGTTTGGAAGAAGAAACTTTCCCATTGGACTTGG
ACATTGCTTACGAAGGTATTATGATTGTTCACAAGGCTTCTGAAGACTCTACTTTCCCAACTAAGATTC
AACACTGTATTAACGAAATTCCAGTTATGTTCTACGAATCTGAATTCGAAATGTTCGAAGCTTTGACTG
ACTTGGTTTTGTTGTTGGACCCAGACATTTTGTCTGGTTTCGAAATTCACAACTTCTCTTGGGGTTACA
TTATTGAAAGATGTCAAAAGATTCACCAATTCGACATTGTTAGAGAATTGGCTAGAGTTAAGTGTCAAA
TTAAGACTAAGTTGTCTGACACTTGGGGTTACGCTCACTCTTCTGGTATTATGATTACTGGTAGACACA
TGATTAACATTTGGAGAGCTTTGAGATCTGACGTTAACTTGACTCAATACACTATTGAATCTGCTGCTT
TCAACATTTTGCACAAGAGATTGCCACACTTCTCTTTCGAATCTTTGACTAACATGTGGAACGCTAAGA
AGTCTACTACTGAATTGAAGACTGTTTTGAACTACTGGTTGTCTAGAGCTCAAATTAACATTCAATTGT
TGAGAAAGCAAGACTACATTGCTAGAAACATTGAACAAGCTAGATTGATTGGTATTGACTTCCACTCTG
TTTACTACAGAGGTTCTCAATTCAAGGTTGAATCTTTCTTGATTAGAATTTGTAAGTCTGAATCTTTCA
TTTTGTTGTCTCCAGGTAAGAAGGACGTTAGAAAGCAAAAGGCTTTGGAATGTGTTCCATTGGTTATGG
AACCAGAATCTGCTTTCTACAAGTCTCCATTGATTGTTTTGGACTTCCAATCTTTGTACCCATCTATTA
TGATTGGTTACAACTACTGTTACTCTACTATGATTGGTAGAGTTAGAGAAATTAACTTGACTGAAAACA
ACTTGGGTGTTTCTAAGTTCTCTTTGCCAAGAAACATTTTGGCTTTGTTGAAGAACGACGTTACTATTG
CTCCAAACGGTGTTGTTTACGCTAAGACTTCTGTTAGAAAGTCTACTTTGTCTAAGATGTTGACTGACA
TTTTGGACGTTAGAGTTATGATTAAGAAGACTATGAACGAAATTGGTGACGACAACACTACTTTGAAGA
GATTGTTGAACAACAAGCAATTGGCTTTGAAGTTGTTGGCTAACGTTACTTACGGTTACACTTCTGCTT
CTTTCTCTGGTAGAATGCCATGTTCTGACTTGGCTGACTCTATTGTTCAAACTGGTAGAGAAACTTTGG
AAAAGGCTATTGACATTATTGAAAAGGACGAAACTTGGAACGCTAAGGTTGTTTACGGTGACACTGACT
CTTTGTTCGTTTACTTGCCAGGTAAGACTGCTATTGAAGCTTTCTCTATTGGTCACGCTATGGCTGAAA
GAGTTACTCAAAACAACCCAAAGCCAATTTTCTTGAAGTTCGAAAAGGTTTACCACCCATCTATTTTGA
TTTCTAAGAAGAGATACGTTGGTTTCTCTTACGAATCTCCATCTCAAACTTTGCCAATTTTCGACGCTA
AGGGTATTGAAACTGTTAGAAGAGACGGTATTCCAGCTCAACAAAAGATTATTGAAAAGTGTATTAGAT
TGTTGTTCCAAACTAAGGACTTGTCTAAGATTAAGAAGTACTTGCAAAACGAATTCTTCAAGATTCAAA

TTGGTAAGGTTTCTGCTCAAGACTTCTGTTTCGCTAAGGAAGTTAAGTTGGGTGCTTACAAGTCTGAAA
AGACTGCTCCAGCTGGTGCTGTTGTTGTTAAGAGAAGAATTAACGAAGACCACAGAGCTGAACCACAAT
ACAAGGAAAGAATTCCATACTTGGTTGTTAAGGGTAAGCAAGGTCAATTGTTGAGAGAAAGATGTGTTT
CTCCAGAAGAATTCTTGGAAGGTGAAAACTTGGAATTGGACTCTGAATACTACATTAACAAGATTTTGA
TTCCACCATTGGACAGATTGTTCAACTTGATTGGTATTAACGTTGGTAACTGGGCTCAAGAAATTGTTA
AGTCTAAGAGAGCTTCTACTACTACTACTAAGGTTGAAAACATTACTAGAGTTGGTACTTCTGCTACTT
GTTGTAACTGTGGTGAAGAATTGACTAAGATTTGTTCTTTGCAATTGTGTGACGACTGTTTGGAAAAGA
GATCTACTACTACTTTGTCTTTCTTGATTAAGAAGTTGAAGAGACAAAAGGAATACCAAACTTTGAAGA
CTGTTTGTAGAACTTGTTCTTACAGATACACTTCTGACGCTGGTATTGAAAACGACCACATTGCTTCTA
AGTGTAACTCTTACGACTGTCCAGTTTTCTACTCTAGAGTTAAGGCTGAAAGATACTTGAGAGACAACC
AATCTGTTCAAAGAGAAGAAGCTTTGATTTCTTTGAACGACTGG**TAA**taaattgaattgaattgaaatc
gatagatcaatttttttcttttctctttccccatcctttacgctaaaataatagtttattttattttt
gaatatttttatttatatacgtatatatagactattatttatctttaatgattattaagatttttat
taaaaaaaaattcgctcctcttttaatgcctttatgcgcggccgc
NotI

*REV3rc-ΔC*

SgrAI
caccggtgtataaaaca**ATG**TCTAGAGAATCTAACGACACTATTCAATCTGACACTGTTAGATCTTCTT
CTAAGTCTGACTACTTCAGAATTCAATTGAACAACCAAGACTACTACATGTCTAAGCCAACTTTCTTGG
ACCCATCTCACGGTGAATCTTTGCCATTGAACCAATTCTCTCAAGTTCCAAACATTAGAGTTTTCGGTG
CTTTGCCAACTGGTCACCAAGTTTTGTGTCACGTTCACGGTATTTTGCCATACATGTTCATTAAGTACG
ACGGTCAAATTACTGACACTTCTACTTTGAGACACCAAAGATGTGCTCAAGTTCACAAGACTTTGGAAG
TTAAGATTAGAGCTTCTTTCAAGAGAAAGAAGGACGACAAGCACGACTTGGCTGGTGACAAGTTGGGTA
ACTTGAACTTCGTTGCTGACGTTTCTGTTGTTAAGGGTATTCCATTCTACGGTTACCACGTTGGTTGGA
ACTTGTTCTACAAGATTTCTTTGTTGAACCCATCTTGTTTGTCTAGAATTTCTGAATTGATTAGAGACG
GTAAGATTTTCGGTAAGAAGTTCGAAATTTACGAATCTCACATTCCATACTTGTTGCAATGGACTGCTG
ACTTCAACTTGTTCGGTTGTTCTTGGATTAACGTTGACAGATGTTACTTCAGATCTCCAGTTTTGAACT
CTATTTTGGACATTGACAAGTTGACTATTAACGACGACTTGCAATTGTTGTTGGACAGATTCTGTGACT
TCAAGTGTAACGTTTTGTCTAGAAGAGACTTCCCAAGAGTTGGTAACGGTTTGATTGAAATTGACATTT
TGCCACAATTCATTAAGAACAGAGAAAAGTTGCAACACAGAGACATTCACCACGACTTCTTGGAAAAGT
TGGGTGACATTTCTGACATTCCAGTTAAGCCATACGTTTCTTCTGCTAGAGACATGATTAACGAATTGA
CTATGCAAAGAGAAGAATTGTCTTTGAAGGAATACAAGGAACCACCAGAAACTAAGAGACACGTTTCTG
GTCACCAATGGCAATCTTCTGGTGAATTCGAAGCTTTCTACAAGAAGGCTCAACACAAGACTTCTACTT
TCGACGGTCAAATTCCAAACTTCGAAAACTTCATTGACAAGAACCAAAAGTTCTCTGCTATTAACACTC
CATACGAAGCTTTGCCACAATTGTGGCCAAGATTGCCACAAATTGAAATTAACAACAACTCTATGCAAG
ACAAGAAGAACGACGACCAAGTTAACGCTTCTTTCACTGAATACGAAATTTGTGGTGTTGACAACGAAA
ACGAAGGTGTTAAGGGTTCTAACATTAAGTCTAGATCTTACTCTTGGTTGCCAGAATCTATTGCTTCTC
CAAAGGACTCTACTATTTTGTTGGACCACCAAACTAAGTACCACAACACTATTAACTTCTCTATGGACT
GTGCTATGACTCAAAACATGGCTTCTAAGAGAAAGTTGAGATCTTCTGTTTCTGCTAACAAGACTTCTT
TGTTGTCTAGAAAGAGAAAGAAGGTTATGGCTGCTGGTTTGAGATACGGTAAGAGAGCTTTCGTTTACG
GTGAACCACCATTCGGTTACCAAGACATTTTGAACAAGTTGGAAGACGAAGGTTTCCCAAAGATTGACT
ACAAGGACCCATTCTTCTCTAACCCAGTTGACTTGGAAAACAAGCCATACGCTTACGCTGGTAAGAGAT
TCGAAATTTCTTCTACTCACGTTTCTACTAGAATTCCAGTTCAATTCGGTGGTGAAACTGTTTCTGTTT
ACAACAAGCCAACTTTCGACATGTTCTCTTCTTGGAAGTACGCTTTGAAGCCACCAACTTACGACGCTG
TTCAAAAGTGGTACAACAAGGTTCCATCTATGGGTAACAAGAAGACTGAATCTCAAATTTCTATGCACA
CTCCACACTCTAAGTTCTTGTACAAGTTCGCTTCTGACGTTTCTGGTAAGCAAAAGAGAAAGAAGTCTT
CTGTTCACGACTCTTTGACTCACTTGACTTTGGAAATTCACGCTAACACTAGATCTGACAAGATTCCAG
ACCCAGCTATTGACGAAGTTTCTATGATTATTTGGTGTTTGGAAGAAGAAACTTTCCCATTGGACTTGG
ACATTGCTTACGAAGGTATTATGATTGTTCACAAGGCTTCTGAAGACTCTACTTTCCCAACTAAGATTC

AACACTGTATTAACGAAATTCCAGTTATGTTCTACGAATCTGAATTCGAAATGTTCGAAGCTTTGACTG
ACTTGGTTTTGTTGTTGGACCCAGACATTTTGTCTGGTTTCGAAATTCACAACTTCTCTTGGGGTTACA
TTATTGAAAGATGTCAAAAGATTCACCAATTCGACATTGTTAGAGAATTGGCTAGAGTTAAGTGTCAAA
TTAAGACTAAGTTGTCTGACACTTGGGGTTACGCTCACTCTTCTGGTATTATGATTACTGGTAGACACA
TGATTAACATTTGGAGAGCTTTGAGATCTGACGTTAACTTGACTCAATACACTATTGAATCTGCTGCTT
TCAACATTTTGCACAAGAGATTGCCACACTTCTCTTTCGAATCTTTGACTAACATGTGGAACGCTAAGA
AGTCTACTACTGAATTGAAGACTGTTTTGAACTACTGGTTGTCTAGAGCTCAAATTAACATTCAATTGT
TGAGAAAGCAAGACTACATTGCTAGAAACATTGAACAAGCTAGATTGATTGGTATTGACTTCCACTCTG
TTTACTACAGAGGTTCTCAATTCAAGGTTGAATCTTTCTTGATTAGAATTTGTAAGTCTGAATCTTTCA
TTTTGTTGTCTCCAGGTAAGAAGGACGTTAGAAAGCAAAAGGCTTTGGAATGTGTTCCATTGGTTATGG
AACCAGAATCTGCTTTCTACAAGTCTCCATTGATTGTTTTGGACTTCCAATCTTTGTACCCATCTATTA
TGATTGGTTACAACTACTGTTACTCTACTATGATTGGTAGAGTTAGAGAAATTAACTTGACTGAAAACA
ACTTGGGTGTTTCTAAGTTCTCTTTGCCAAGAAACATTTTGGCTTTGTTGAAGAACGACGTTACTATTG
CTCCAAACGGTGTTGTTTACGCTAAGACTTCTGTTAGAAAGTCTACTTTGTCTAAGATGTTGACTGACA
TTTTGGACGTTAGAGTTATGATTAAGAAGACTATGAACGAAATTGGTGACGACAACACTACTTTGAAGA
GATTGTTGAACAACAAGCAATTGGCTTTGAAGTTGTTGGCTAACGTTACTTACGGTTACACTTCTGCTT
CTTTCTCTGGTAGAATGCCATGTTCTGACTTGGCTGACTCTATTGTTCAAACTGGTAGAGAAACTTTGG
AAAAGGCTATTGACATTATTGAAAAGGACGAAACTTGGAACGCTAAGGTTGTTTACGGTGACACTGACT
CTTTGTTCGTTTACTTGCCAGGTAAGACTGCTATTGAAGCTTTCTCTATTGGTCACGCTATGGCTGAAA
GAGTTACTCAAAACAACCCAAAGCCAATTTTCTTGAAGTTCGAAAAGGTTTACCACCCATCTATTTTGA
TTTCTAAGAAGAGATACGTTGGTTTCTCTTACGAATCTCCATCTCAAACTTTGCCAATTTTCGACGCTA
AGGGTATTGAAACTGTTAGAAGAGACGGTATTCCAGCTCAACAAAAGATTATTGAAAAGTGTATTAGAT
TGTTGTTCCAAACTAAGGACTTGTCTAAGATTAAGAAGTACTTGCAAAACGAATTCTTCAAGATTCAAA
TTGGTAAGGTTTCTGCTCAAGACTTCTGTTTCGCTAAGGAAGTTAAGTTGGGTGCTTACAAGTCTGAAA
AGACTGCTCCAGCTGGTGCTGTTGTTGTTAAGAGAAGAATTAACGAAGACCACAGAGCTGAACCACAAT
ACAAGGAAAGAATTCCATACTTGGTTGTTAAGGGTAAGCAAGGTCAATTGTTGAGAGAAAGATGTGTTT
CTCCAGAAGAATTCTTGGAAGGTGAAAACTTGGAATTGGACTCTGAATACTACATTAACAAGATTTTGA
TTCCACCATTGGACAGATTGTTCAACTTGATTGGTATTAACGTTGGTAACTGGGCTCAAGAAATTGTTA
AGTCTAAGAGAGCTTCT**TAA**taaattgaattgaattgaaatcgatagatcaatttttttcttttctctt
tccccatcctttacgctaaaataatagtttattttatttttttgaatattttttatttatatacgtatat
atagactattatttatcttttaatgattattaagatttttattaaaaaaaaattcgctcctcttttaat
gcctttatgcgcggccgc
NotI

*REV3rc-ΔCh6*

SgrAI
caccggtgtataaaaca**ATG**TCTAGAGAATCTAACGACACTATTCAATCTGACACTGTTAGATCTTCTT
CTAAGTCTGACTACTTCAGAATTCAATTGAACAACCAAGACTACTACATGTCTAAGCCAACTTTCTTGG
ACCCATCTCACGGTGAATCTTTGCCATTGAACCAATTCTCTCAAGTTCCAAACATTAGAGTTTTCGGTG
CTTTGCCAACTGGTCACCAAGTTTTGTGTCACGTTCACGGTATTTTGCCATACATGTTCATTAAGTACG
ACGGTCAAATTACTGACACTTCTACTTTGAGACACCAAAGATGTGCTCAAGTTCACAAGACTTTGGAAG
TTAAGATTAGAGCTTCTTTCAAGAGAAAGAAGGACGACAAGCACGACTTGGCTGGTGACAAGTTGGGTA
ACTTGAACTTCGTTGCTGACGTTTCTGTTGTTAAGGGTATTCCATTCTACGGTTACCACGTTGGTTGGA
ACTTGTTCTACAAGATTTCTTTGTTGAACCCATCTTGTTTGTCTAGAATTTCTGAATTGATTAGAGACG
GTAAGATTTTCGGTAAGAAGTTCGAAATTTACGAATCTCACATTCCATACTTGTTGCAATGGACTGCTG
ACTTCAACTTGTTCGGTTGTTCTTGGATTAACGTTGACAGATGTTACTTCAGATCTCCAGTTTTGAACT
CTATTTTGGACATTGACAAGTTGACTATTAACGACGACTTGCAATTGTTGTTGGACAGATTCTGTGACT
TCAAGTGTAACGTTTTGTCTAGAAGAGACTTCCCAAGAGTTGGTAACGGTTTGATTGAAATTGACATTT

TGCCACAATTCATTAAGAACAGAGAAAAGTTGCAACACAGAGACATTCACCACGACTTCTTGGAAAAGT
TGGGTGACATTTCTGACATTCCAGTTAAGCCATACGTTTCTTCTGCTAGAGACATGATTAACGAATTGA
CTATGCAAAGAGAAGAATTGTCTTTGAAGGAATACAAGGAACCACCAGAAACTAAGAGACACGTTTCTG
GTCACCAATGGCAATCTTCTGGTGAATTCGAAGCTTTCTACAAGAAGGCTCAACACAAGACTTCTACTT
TCGACGGTCAAATTCCAAACTTCGAAAACTTCATTGACAAGAACCAAAAGTTCTCTGCTATTAACACTC
CATACGAAGCTTTGCCACAATTGTGGCCAAGATTGCCACAAATTGAAATTAACAACAACTCTATGCAAG
ACAAGAAGAACGACGACCAAGTTAACGCTTCTTTCACTGAATACGAAATTTGTGGTGTTGACAACGAAA
ACGAAGGTGTTAAGGGTTCTAACATTAAGTCTAGATCTTACTCTTGGTTGCCAGAATCTATTGCTTCTC
CAAAGGACTCTACTATTTTGTTGGACCACCAAACTAAGTACCACAACACTATTAACTTCTCTATGGACT
GTGCTATGACTCAAAACATGGCTTCTAAGAGAAAGTTGAGATCTTCTGTTTCTGCTAACAAGACTTCTT
TGTTGTCTAGAAAGAGAAAGAAGGTTATGGCTGCTGGTTTGAGATACGGTAAGAGAGCTTTCGTTTACG
GTGAACCACCATTCGGTTACCAAGACATTTTGAACAAGTTGGAAGACGAAGGTTTCCCAAAGATTGACT
ACAAGGACCCATTCTTCTCTAACCCAGTTGACTTGGAAAACAAGCCATACGCTTACGCTGGTAAGAGAT
TCGAAATTTCTTCTACTCACGTTTCTACTAGAATTCCAGTTCAATTCGGTGGTGAAACTGTTTCTGTTT
ACAACAAGCCAACTTTCGACATGTTCTCTTCTTGGAAGTACGCTTTGAAGCCACCAACTTACGACGCTG
TTCAAAAGTGGTACAACAAGGTTCCATCTATGGGTAACAAGAAGACTGAATCTCAAATTTCTATGCACA
CTCCACACTCTAAGTTCTTGTACAAGTTCGCTTCTGACGTTTCTGGTAAGCAAAAGAGAAAGAAGTCTT
CTGTTCACGACTCTTTGACTCACTTGACTTTGGAAATTCACGCTAACACTAGATCTGACAAGATTCCAG
ACCCAGCTATTGACGAAGTTTCTATGATTATTTGGTGTTTGGAAGAAGAAACTTTCCCATTGGACTTGG
ACATTGCTTACGAAGGTATTATGATTGTTCACAAGGCTTCTGAAGACTCTACTTTCCCAACTAAGATTC
AACACTGTATTAACGAAATTCCAGTTATGTTCTACGAATCTGAATTCGAAATGTTCGAAGCTTTGACTG
ACTTGGTTTTGTTGTTGGACCCAGACATTTTGTCTGGTTTCGAAATTCACAACTTCTCTTGGGGTTACA
TTATTGAAAGATGTCAAAAGATTCACCAATTCGACATTGTTAGAGAATTGGCTAGAGTTAAGTGTCAAA
TTAAGACTAAGTTGTCTGACACTTGGGGTTACGCTCACTCTTCTGGTATTATGATTACTGGTAGACACA
TGATTAACATTTGGAGAGCTTTGAGATCTGACGTTAACTTGACTCAATACACTATTGAATCTGCTGCTT
TCAACATTTTGCACAAGAGATTGCCACACTTCTCTTTCGAATCTTTGACTAACATGTGGAACGCTAAGA
AGTCTACTACTGAATTGAAGACTGTTTTGAACTACTGGTTGTCTAGAGCTCAAATTAACATTCAATTGT
TGAGAAAGCAAGACTACATTGCTAGAAACATTGAACAAGCTAGATTGATTGGTATTGACTTCCACTCTG
TTTACTACAGAGGTTCTCAATTCAAGGTTGAATCTTTCTTGATTAGAATTTGTAAGTCTGAATCTTTCA
TTTTGTTGTCTCCAGGTAAGAAGGACGTTAGAAAGCAAAAGGCTTTGGAATGTGTTCCATTGGTTATGG
AACCAGAATCTGCTTTCTACAAGTCTCCATTGATTGTTTTGGACTTCCAATCTTTGTACCCATCTATTA
TGATTGGTTACAACTACTGTTACTCTACTATGATTGGTAGAGTTAGAGAAATTAACTTGACTGAAAACA
ACTTGGGTGTTTCTAAGTTCTCTTTGCCAAGAAACATTTTGGCTTTGTTGAAGAACGACGTTACTATTG
CTCCAAACGGTGTTGTTTACGCTAAGACTTCTGTTAGAAAGTCTACTTTGTCTAAGATGTTGACTGACA
TTTTGGACGTTAGAGTTATGATTAAGAAGACTATGAACGAAATTGGTGACGACAACACTACTTTGAAGA
GATTGTTGAACAACAAGCAATTGGCTTTGAAGTTGTTGGCTAACGTTACTTACGGTTACACTTCTGCTT
CTTTCTCTGGTAGAATGCCATGTTCTGACTTGGCTGACTCTATTGTTCAAACTGGTAGAGAAACTTTGG
AAAAGGCTATTGACATTATTGAAAAGGACGAAACTTGGAACGCTAAGGTTGTTTACGGTGACACTGACT
CTTTGTTCGTTTACTTGCCAGGTAAGACTGCTATTGAAGCTTTCTCTATTGGTCACGCTATGGCTGAAA
GAGTTACTCAAAACAACCCAAAGCCAATTTTCTTGAAGTTCGAAAAGGTTTACCACCCATCTATTTTGA
TTTCTAAGAAGAGATACGTTGGTTTCTCTTACGAATCTCCATCTCAAACTTTGCCAATTTTCGACGCTA
AGGGTATTGAAACTGTTAGAAGAGACGGTATTCCAGCTCAACAAAAGATTATTGAAAAGTGTATTAGAT
TGTTGTTCCAAACTAAGGACTTGTCTAAGATTAAGAAGTACTTGCAAAACGAATTCTTCAAGATTCAAA
TTGGTAAGGTTTCTGCTCAAGACTTCTGTTTCGCTAAGGAAGTTAAGTTGGGTGCTTACAAGTCTGAAA
AGACTGCTCCAGCTGGTGCTGTTGTTGTTAAGAGAAGAATTAACGAAGACCACAGAGCTGAACCACAAT
ACAAGGAAAGAATTCCATACTTGGTTGTTAAGGGTAAGCAAGGTCAATTGTTGAGAGAAAGATGTGTTT
CTCCAGAAGAATTCTTGGAAGGTGAAAACTTGGAATTGGACTCTGAATACTACATTAACAAGATTTTGA
TTCCACCATTGGACAGATTGTTCAACTTGATTGGTATTAACGTTGGTAACTGGGCTCAAGAAATTGTTA
AGTCTAAGAGAGCTTCTCATCACCACCATCACCAT**TAA**taaattgaattgaattgaaatcgatagatca

atttttttcttttctctttccccatcctttacgctaaaataatagtttatttatttttttgaatatttt
ttatttatatacgtatatatagactattatttatcttttaatgattattaagattttattaaaaaaaa
attcgctcctcttttaatgcctttatgcgcggccgc
NotI

*REV7rc* AscI - fragment:

AscI
ggcgcgcctataaaaca**ATG**AACAGATGGGTTGAAAAGTGGTTGAGAGTTTACTTGAAGTGTTACATTA
ACTTGATTTTGTTCTACAGAAACGTTTACCCACCACAATCTTTCGACTACACTACTTACCAATCTTTCA
ACTTGCCACAATTCGTTCCAATTAACAGACACCCAGCTTTGATTGACTACATTGAAGAATTGATTTTGG
ACGTTTTGTCTAAGTTGACTCACGTTTACAGATTCTCTATTTGTATTATTAACAAGAAGAACGACTTGT
GTATTGAAAAGTACGTTTTGGACTTCTCTGAATTGCAACACGTTGACAAGGACGACCAAATTATTACTG
AAACTGAAGTTTTCGACGAATTCAGATCTTCTTTGAACTCTTTGATTATGCACTTGGAAAAGTTGCCAA
AGGTTAACGACGACACTATTACTTTCGAAGCTGTTATTAACGCTATTGAATTGGAATTGGGTCACAAGT
TGGACAGAAACAGAAGAGTTGACTCTTTGGAAGAAAAGGCTGAAATTGAAAGAGACTCTAACTGGGTTA
AGTGTCAAGAAGACGAAAACTTGCCAGACAACAACGGTTTCCAACCACCAAAGATTAAGTTGACTTCTT
TGGTTGGTTCTGACGTTGGTCCATTGATTATTCACCAATTCTCTGAAAAGTTGATTTCTGGTGACGACA
AGATTTTGAACGGTGTTTACTCTCAATACGAAGAAGGTGAATCTATTTTCGGTTCTTTGTTC**TAA**taaa
ttgaattgaattgaaatcgatagatcaattttttttcttttctctttccccatcctttacgctaaaataa
tagtttatttatttttttgaatattttttatttatatacgtatatatagactattatttatcttttaat
gattattaagattttattaaaaaaaaattcgctcctcttttaatgcctttatgcctcgag
XhoI

*REV1rc* SgrAI – NotI fragment:

SgrAI
caccggtgtataaaaca**ATG**GGTGAACATGGTGGTTTGGTTGATTTGTTGGATTCCGATTTGGAGTACT
CTATCAACAGAGAAACTCCAGATAAGAACAACTGCTTGTCCCAACAATCTGTTAACGATTCTCATTTGA
CTGCTAAAACCGGTGGTTTGAATGCTAGATCTTTCTTGTCTACCTTGTCCGATGATTCCTTGATCGAAT
ACGTCAATCAATTGTCTCAGACCAACAAGAACAATTCTAATCCAACTGCTGGTACTTTGAGATTCACCA
CTAAGAACATCTCTTGCGACGAATTGCATGCTGACTTAGGTGGTGGTGAAGATTCTCCAATTGCTAGAT
CCGTTATCGAAATCCAAGAATCCGATTCTAATGGTGATGACGTTAAGAAGAACACCGTTTACACTAGAG
AAGCTTACTTTCACGAAAAGGCTCATGGTCAAACATTGCAAGACCAAATTTTGAAGGATCAGTACAAGG
ACCAGATCTCCTCTCAATCTTCCAAAATTTTCAAGAACTGCGTCATCTACATCAACGGTTATACAAAAC
CAGGTAGATTGCAGTTGCACGAAATGATCGTTTTACACGGTGGTAAGTTCTTGCACTACCTGTCCTCTA
AAAAGACTGTTACTCATATCGTTGCCTCCAACTTGCCATTGAAGAAGAGAATTGAATTCGCCAACTACA
AGGTTGTTTCTCCAGATTGGATAGTCGACTCTGTAAAAGAGGCTAGATTATTGCCTTGGCAGAACTACT
CTTTGACCTCTAAATTGGACGAACAGCAAAAGAAGTTGGATAACTGTAAGACCGTTAACTCCATTCCTT
TGCCATCTGAAACTTCATTGCATAAGGGTTCTAAGTGTGTTGGTTCTGCTTTGTTGCCAGTTGAACAAC
AATCTCCAGTTAACTTGAACAACTTGGAAGCTAAAAGAATCGTCGCTTGTGATGATCCAGATTTCTTGA
CTTCTTACTTCGCCCATTCTAGATTGCATCATTTGTCTGCTTGGAAGGCTAACTTGAAGGACAAGTTTT
TGAACGAGAACATCCACAAGTACACCAAGATTACTGATAAGGACACCTACATCATCTTCCACATTGATT
TCGATTGCTTTTTCGCTACCGTTGCTTACTTGTGTAGATCCTCTTCTTTTTCTGCTTGCGACTTTAAGA
GAGATCCAATCGTTGTTTGTCACGGCACTAAGAATTCTGATATTGCTTCTTGTAACTACGTCGCCAGAT
CTTACGGTATTAAGAATGGTATGTGGGTTTCCCAAGCTGAAAAGATGTTGCCAAATGGTATCAAGTTGA
TCTCTTTGCCATACACCTTCGAACAGTTCCAATTGAAGTCTGAAGCTTTCTACTCTACCTTGAAGAGGC
TGAACATCTTCAATTTGATCCTGCCAATCTCCATTGATGAAGCTGTTTGCGTTAGAATCATCCCAGATA
ACATTCATAACACCAACACCTTGAACGCTAGATTGTGCGAAGAAATTAGACAAGAGATCTTCCAAGGTA
CTAATGGTTGCACTGTTTCTATTGGTTGCTCCGATTCTTTGGTTTTGGCTAGATTGGCTTTGAAAATGG
CTAAGCCAAACGGTTACAACATCACCTTTAAGTCCAACTTGTCCGAAGAGTTTTGGTCCTCTTTCAAAT

TGGATGATTTGCCTGGTGTTGGTCACTCTACTTTGTCTAGATTAGAATCCACTTTCGACTCCCCACATT
CTTTGAATGATTTGAGAAAAAGGTACACCTTGGATGCTTTGAAAGCTTCCGTTGGTTCTAAACTGGGTA
TGAAGATTCATTTGGCATTGCAAGGTCAAGACGACGAAGAATCATTGAAGATCTTGTACGACCCTAAAG
AAGTCTTGCAGAGAAAGTCTTTGTCCATCGATATTAACTGGGGCATCAGATTCAAGAACATCACCCAAG
TTGACTTGTTCATCGAAAGAGGTTGTCAGTACTTGTTGGAAAAGCTGAACGAAATCAACAAGACTACCT
CTCAGATCACCTTGAAGTTGATGAGAAGATGTAAGGATGCTCCTATTGAACCACCAAAGTATATGGGTA
TGGGTAGATGTGATTCGTTCTCTAGATCATCAAGATTGGGTATCCCAACTAACGAGTTCGGTATTATTG
CTACCGAAATGAAGTCCCTGTATAGAACTTTGGGTTGTCCACCAATGGAATTGAGAGGTTTGGCTTTAC
AATTCAACAAGTTGGTTGATGTTGGCCCAGACAACAATCAGTTGAAATTGAGATTGCCATTCAAGACCA
TCGTTACCAATAGAGCTTTTGAAGCTTTGCCTGAAGATGTCAAGAACGACATTAACAACGAATTCGAGA
AGCGTAACTACAAGAGGAAAGAATCTGGTTTGACCTCCAACTCTTTGTCCTCAAAGAAAAAGGGTTTCG
CCATTTCTAGGTTGGAAGTTAATGACTTGCCATCCACTATGGAAGAACAGTTCATGAATGAATTGCCAA
CGCAAATTAGAGCCGAAGTTAGACATGACTTGAGAATCCAAAAGAAGATCCAACAGACTAAGTTGGGTA
ACTTGCAAGAAAAGATCAAGAGGCGTGAAGAGTCATTGCAAAACGAAAAGAACCATTTCATGGGCCAGA
ACTCTATTTTTCAGCCAATCAAGTTCCAAAACCTGACCAGGTTTAAGAAGATCTGCCAATTGGTTAAGC
AATGGGTTGCTGAAACTTTAGGTGATGGTGGTCCACACGAAAAAGACGTTAAGTTGTTTGTGAAGTACC
TGATCAAGTTGTGCGATTCCAACAGAGTTCATTTGGTGTTGCATCTGTCAAACTTGATCTCTAGGGAAT
TGAACTTGTGCGCTTTTTTGAACCAGGATCATTCTGGTTTTCAAACCTGGGAGAGAATCTTGTTGAACG
ATATTATCCCTCTGCTGAACAGAAACAAACATACCTACCAAACCGTTCGTAAGTTGGACATGGATTTTG
AAGTG**TAA**attgaattgaattgaaatcgatagatcaatttttttcttttctctttccccatcctttacg
ctaaaataatagtttattttatttttttgaatatttttttatttatatacgtatatatagactattattta
tcttttaatgattattaagatttttattaaaaaaaaattcgctcctcttttaatgcctttatgcgcggc
cgc
NotI

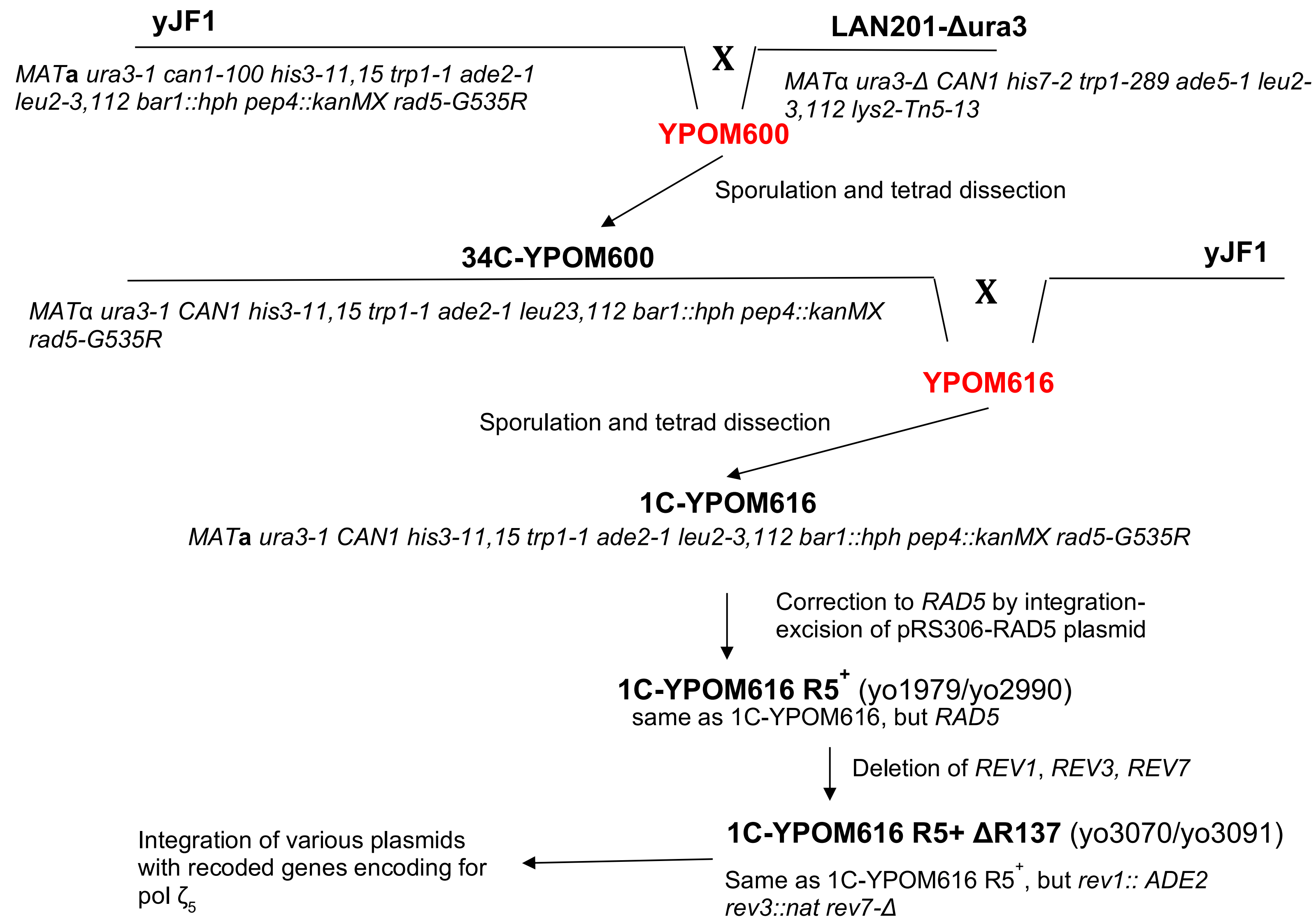

Supplemental Fig 2. Genealogy of the basic yeast strains.
Standard names are in bold and numbers in the laboratory collection are in parenthesis (yo…)

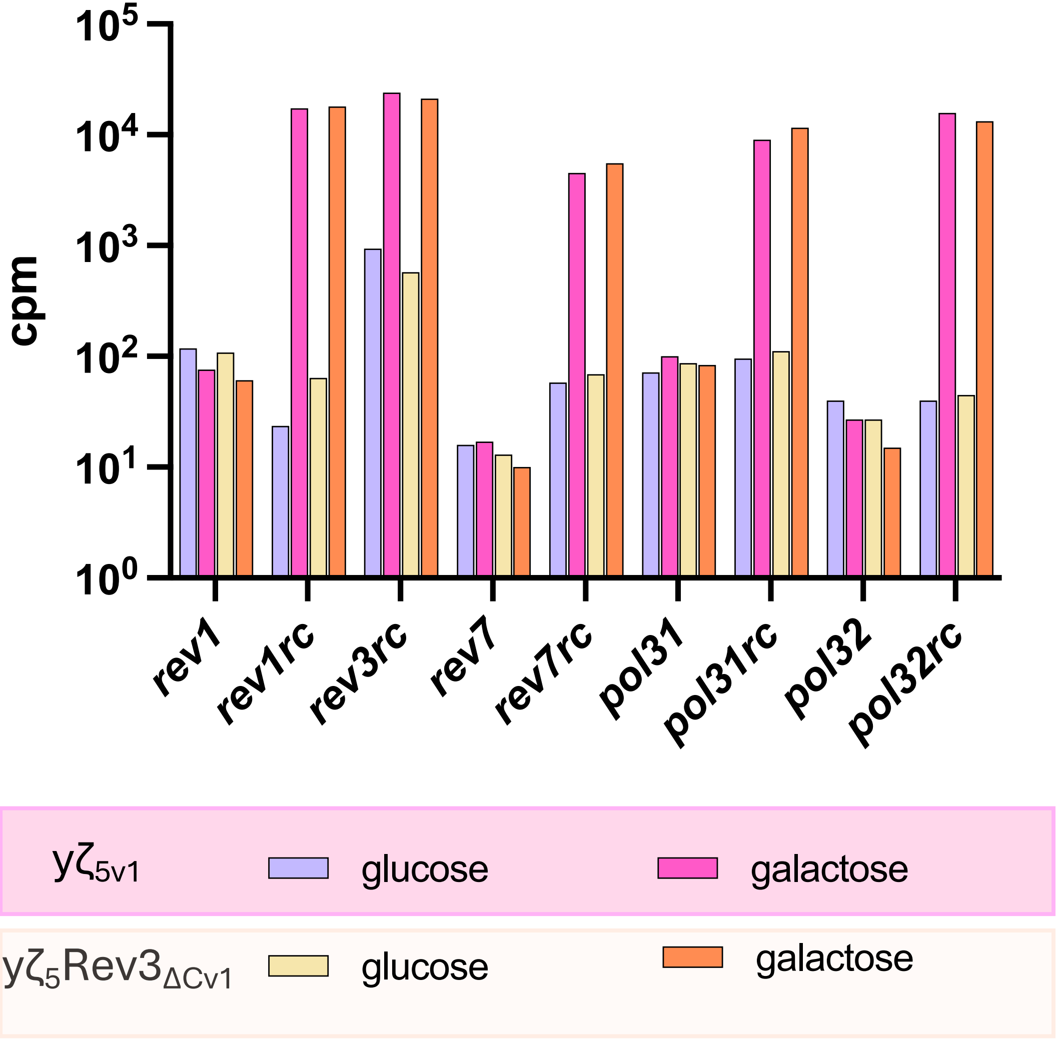

Supplemental Fig 3. Similar patterns of gene expression in strains with five–subunit pol ζ and its CTD-less variant without or with galactose promoter induction.

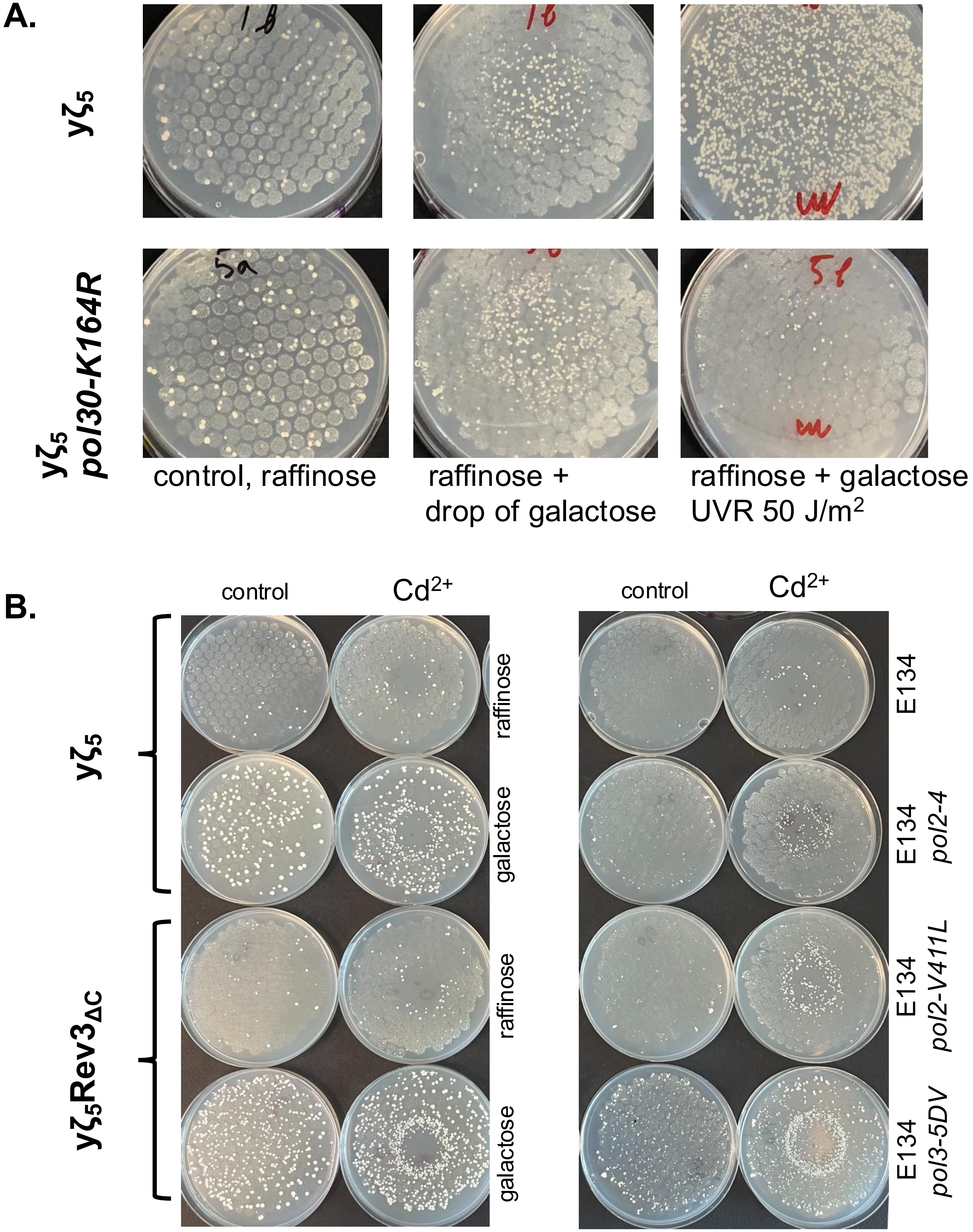

sFig. 4. Pol ζ induces mutations during replication. A. Inability to ubiquitylate PCNA abolishes UVR-induced mutagenesis but does not affect spontaneous mutator effect of elevated pol $ζ_5$. B. Inhibition of MMR by $Cd2^+$ increases mutator effect of pol ζ expression like in replicative pols mutants.

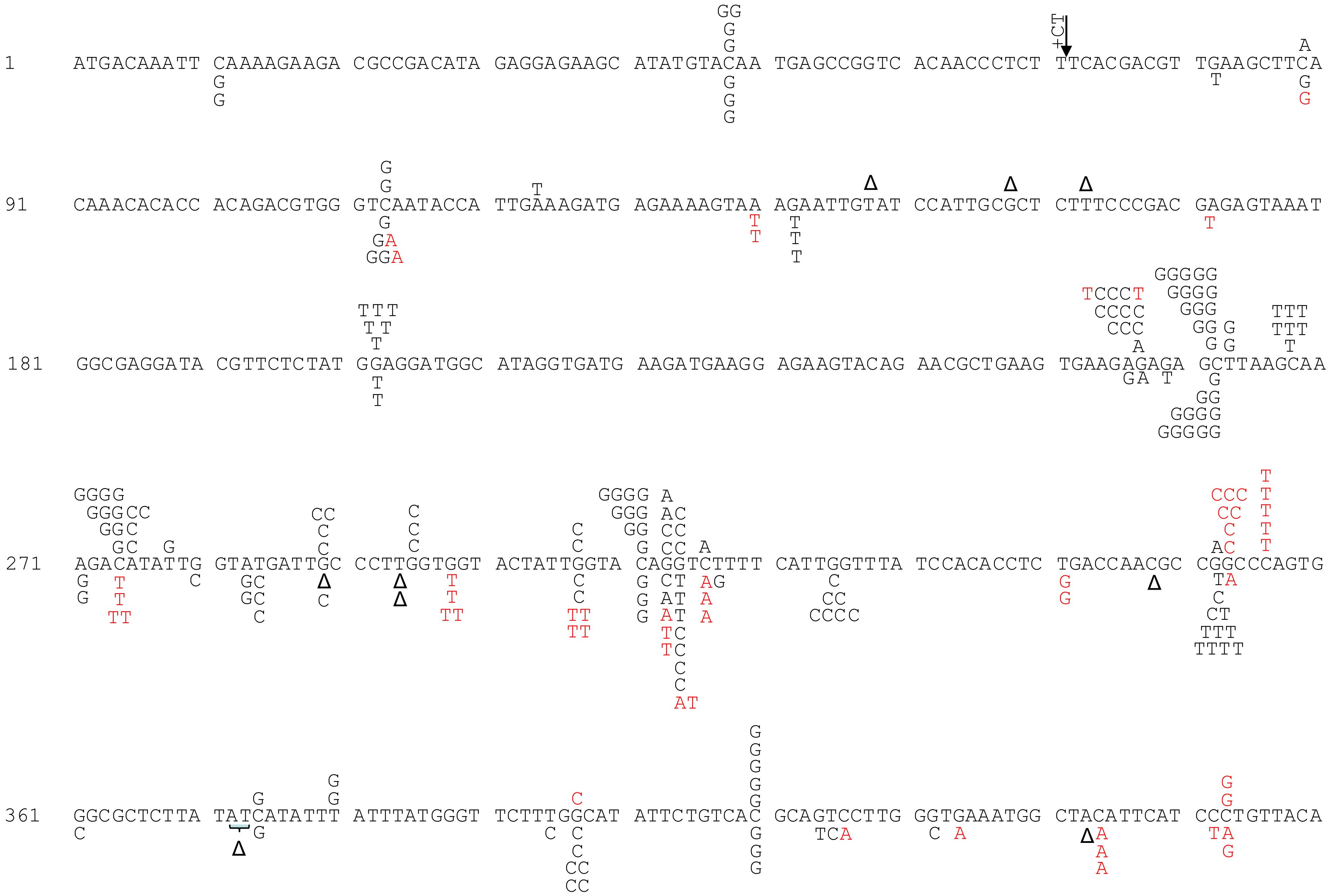

+CT
1 ATGACAAATT CAAAAGAAGA CGCCGACATA GAGGAGAAGC ATATGTACAA TGAGCCGGTC ACAACCCTCT TTCACGACGT TGAAGCTTCA
91 CAAACACACC ACAGACGTGG GTCAATACCA TTGAAAGATG AGAAAAGTAA AGAATTGTAT CCATTGCGCT CTTTCCCGAC GAGAGTAAAT
181 GGCGAGGATA CGTTCTCTAT GGAGGATGGC ATAGGTGATG AAGATGAAGG AGAAGTACAG AACGCTGAAG TGAAGAGAGA GCTTAAGCAA
271 AGACATATTG GTATGATTGC CCTTGGTGGT ACTATTGGTA CAGGTCTTTT CATTGGTTTA TCCACACCTC TGACCAACGC CGGCCCAGTG
361 GGCGCTCTTA TATCATATTT ATTTATGGGT TCTTTGGCAT ATTCTGTCAC GCAGTCCTTG GGTGAAATGG CTACATTCAT CCCTGTTACA

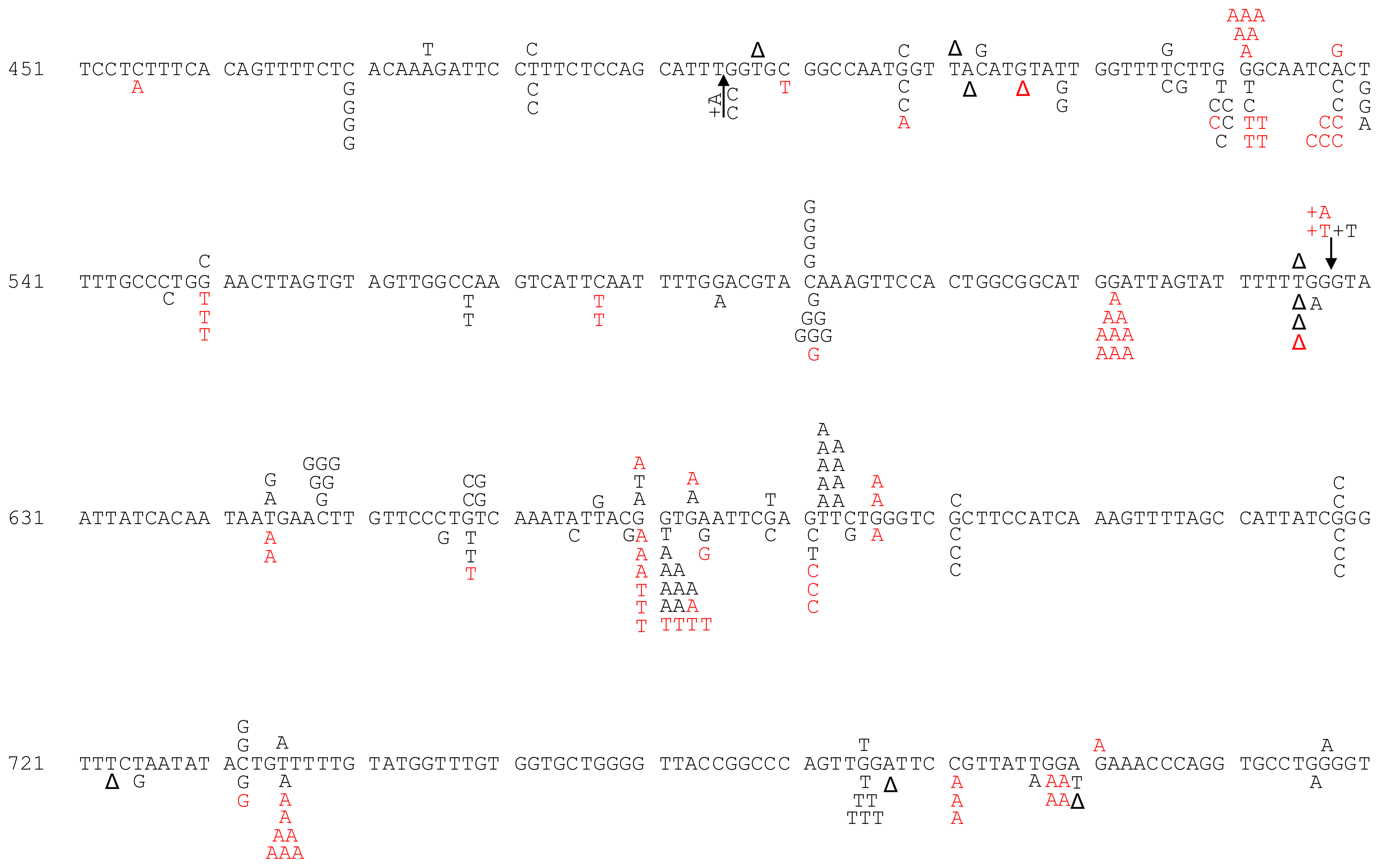
451 TCCTCTTTCA CAGTTTTCTC ACAAAGATTC CTTTCTCCAG CATTTGGTGC GGCCAATGGT TACATGTATT GGTTTTCTTG GGCAATCACT
541 TTTGCCCTGG AACTTAGTGT AGTTGGCCAA GTCATTCAAT TTTGGACGTA CAAAGTTCCA CTGGCGGCAT GGATTAGTAT TTTTTGGGTA
631 ATTATCACAA TAATGAACTT GTTCCCTGTC AAATATTACG GTGAATTCGA GTTCTGGGTC GCTTCCATCA AAGTTTTAGC CATTATCGGG
721 TTTCTAATAT ACTGTTTTTG TATGGTTTGT GGTGCTGGGG TTACCGGCCC AGTTGGATTC CGTTATTGGA GAAACCCAGG TGCCTGGGGT

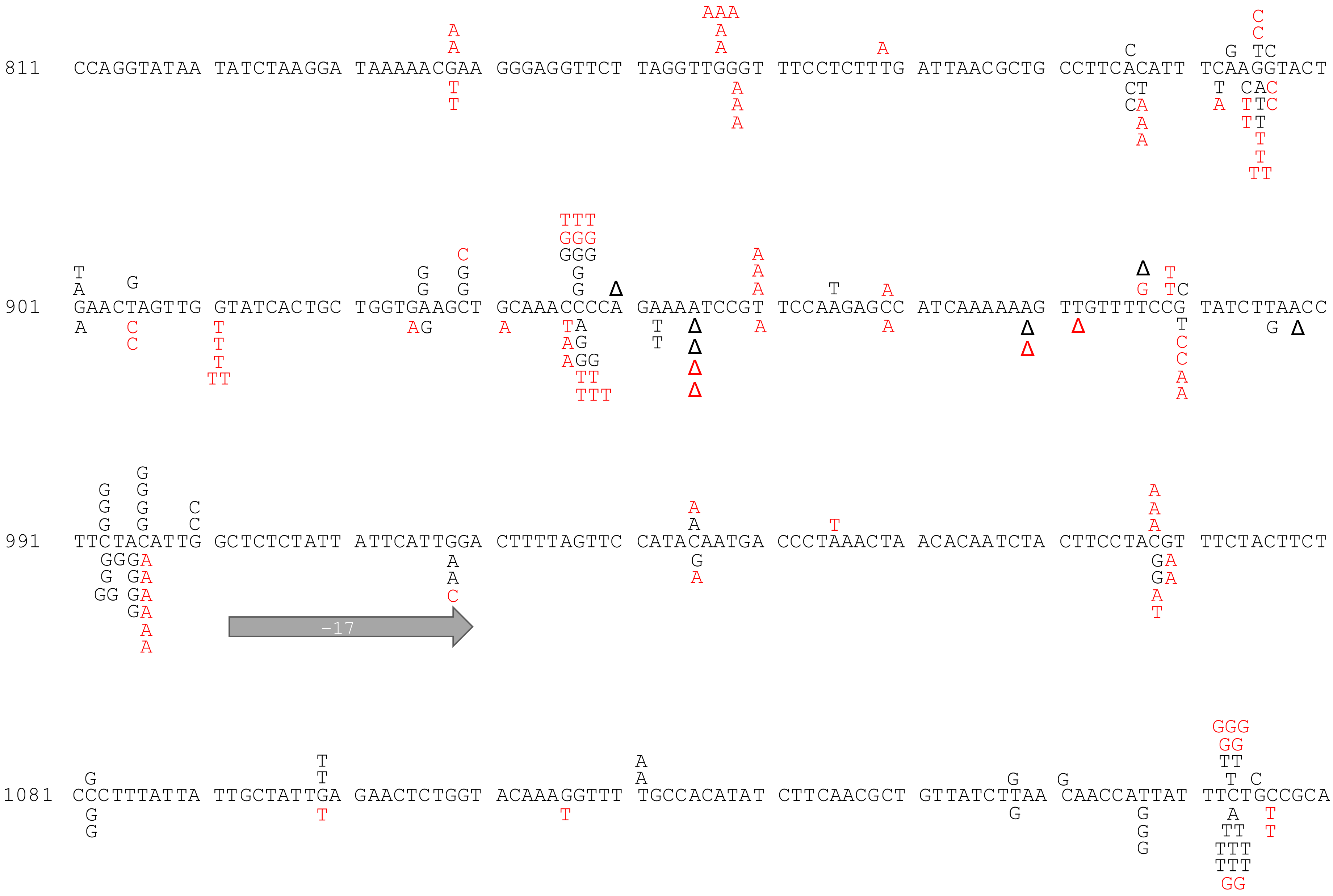
811 CCAGGTATAA TATCTAAGGA TAAAAACGAA GGGAGGTTCT TAGGTTGGGT TTCCTCTTTG ATTAACGCTG CCTTCACATT TCAAGGTACT
901 GAACTAGTTG GTATCACTGC TGGTGAAGCT GCAAACCCCA GAAAATCCGT TCCAAGAGCC ATCAAAAAAG TTGTTTTCCG TATCTTAACC
991 TTCTACATTG GCTCTCTATT ATTCATTGGA CTTTTAGTTC CATACAATGA CCCTAAACTA ACACAATCTA CTTCCTACGT TTCTACTTCT
-17
1081 CCCTTTATTA TTGCTATTGA GAACTCTGGT ACAAAGGTTT TGCCACATAT CTTCAACGCT GTTATCTTAA CAACCATTAT TTCTGCCGCA

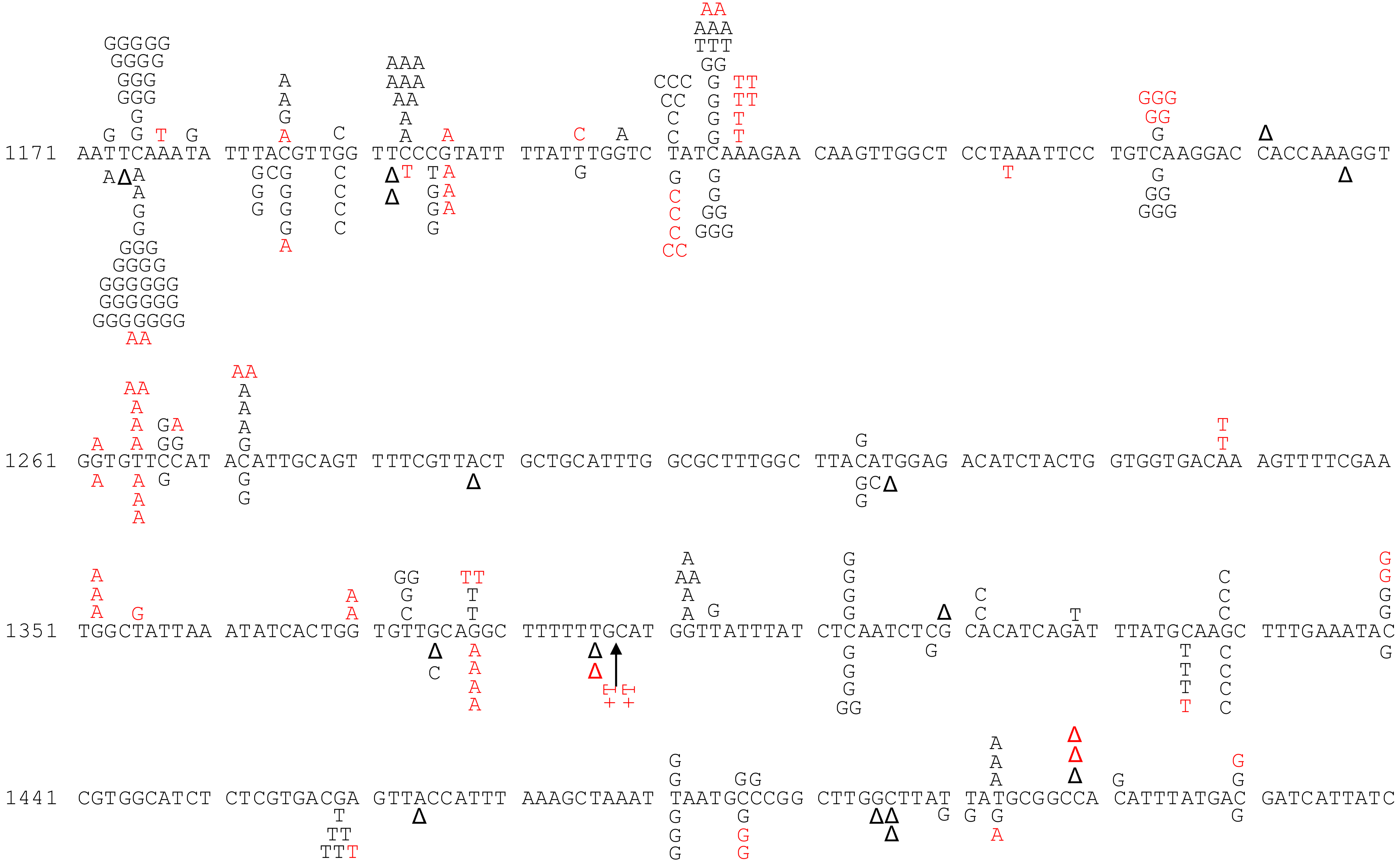
1171 AATTCAAATA TTTACGTTGG TTCCCGTATT TTATTTGGTC TATCAAAGAA CAAGTTGGCT CCTAAATTCC TGTCAAGGAC CACCAAAGGT
1261 GGTGTTCCAT ACATTGCAGT TTTCGTTACT GCTGCATTTG GCGCTTTGGC TTACATGGAG ACATCTACTG GTGGTGACAA AGTTTTCGAA
1351 TGGCTATTAA ATATCACTGG TGTTGCAGGC TTTTTTGCAT GGTTATTTAT CTCAATCTCG CACATCAGAT TTATGCAAGC TTTGAAATAC
1441 CGTGGCATCT CTCGTGACGA GTTACCATTT AAAGCTAAAT TAATGCCCGG CTTGGCTTAT TATGCGGCCA CATTTATGAC GATCATTATC

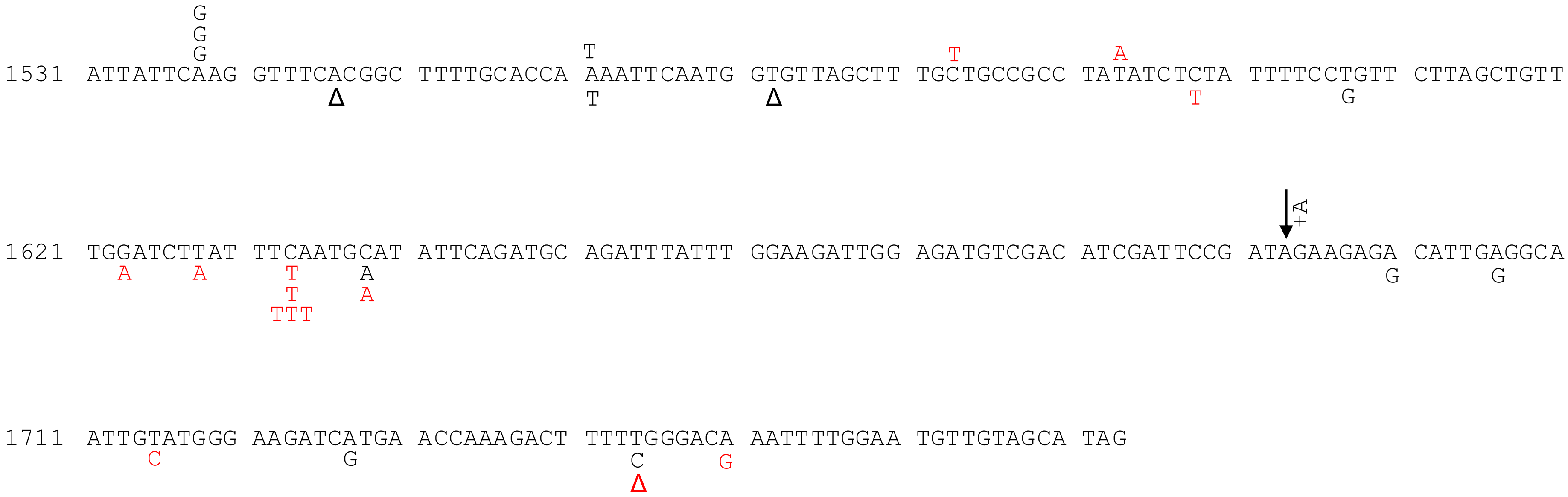

Supplemental Fig. 5. **Mutation spectra produced by pol ζ mutasome variants**

Black letters/symbols - changes found in y$\zeta_5$ strains are above the wild-type DNA sequence, and changes in y$\zeta_5$Rev3$_{\Delta C}$ are under it. Changes in corresponding strains possessing Rev1$_{cat}$ are in red. Points of Insertions are represented by arrows with types of nucleotides inserted. Δ stands for a deletion and a large arrow stand for a larger deletion.

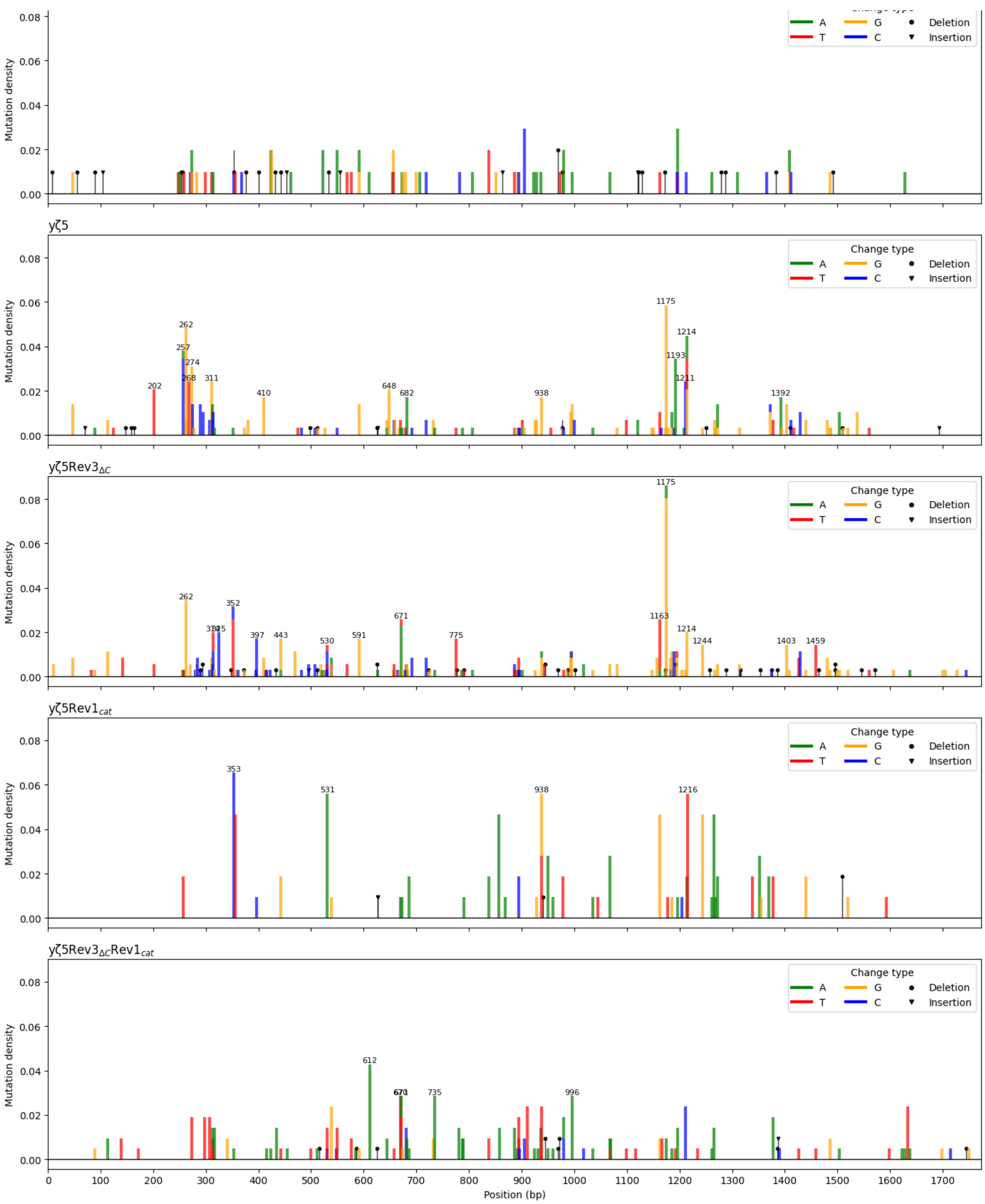

Supplemental Figure 6. Mutation density in strain with elevated pol ζ exhibits strong mutation hotspots as opposed to spectrum of spontaneous mutations in a wild-type strain..

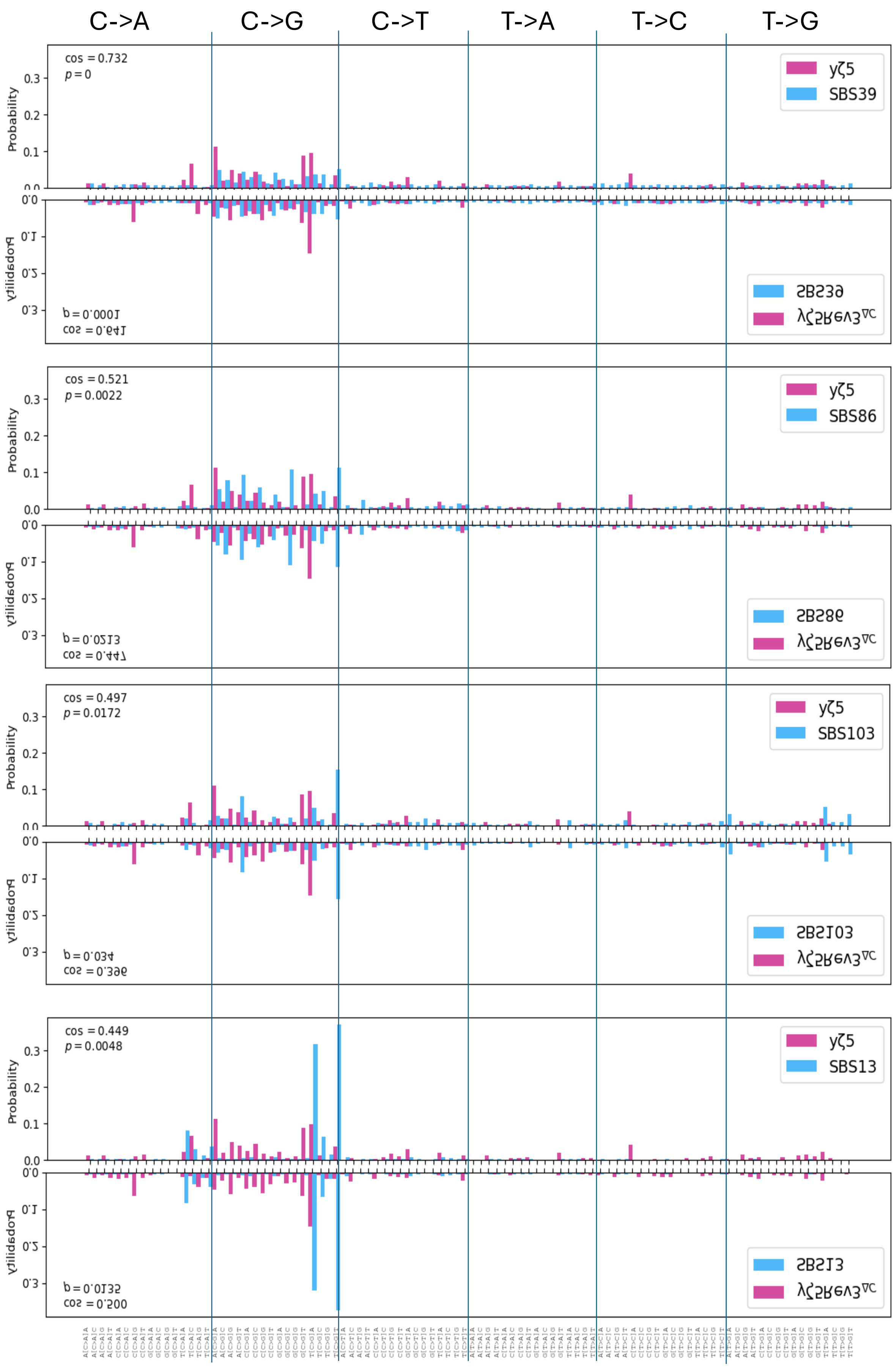

Supplemental Figure 7. Spectra generated by elevated pol ζ mirror cancer mutation signatures

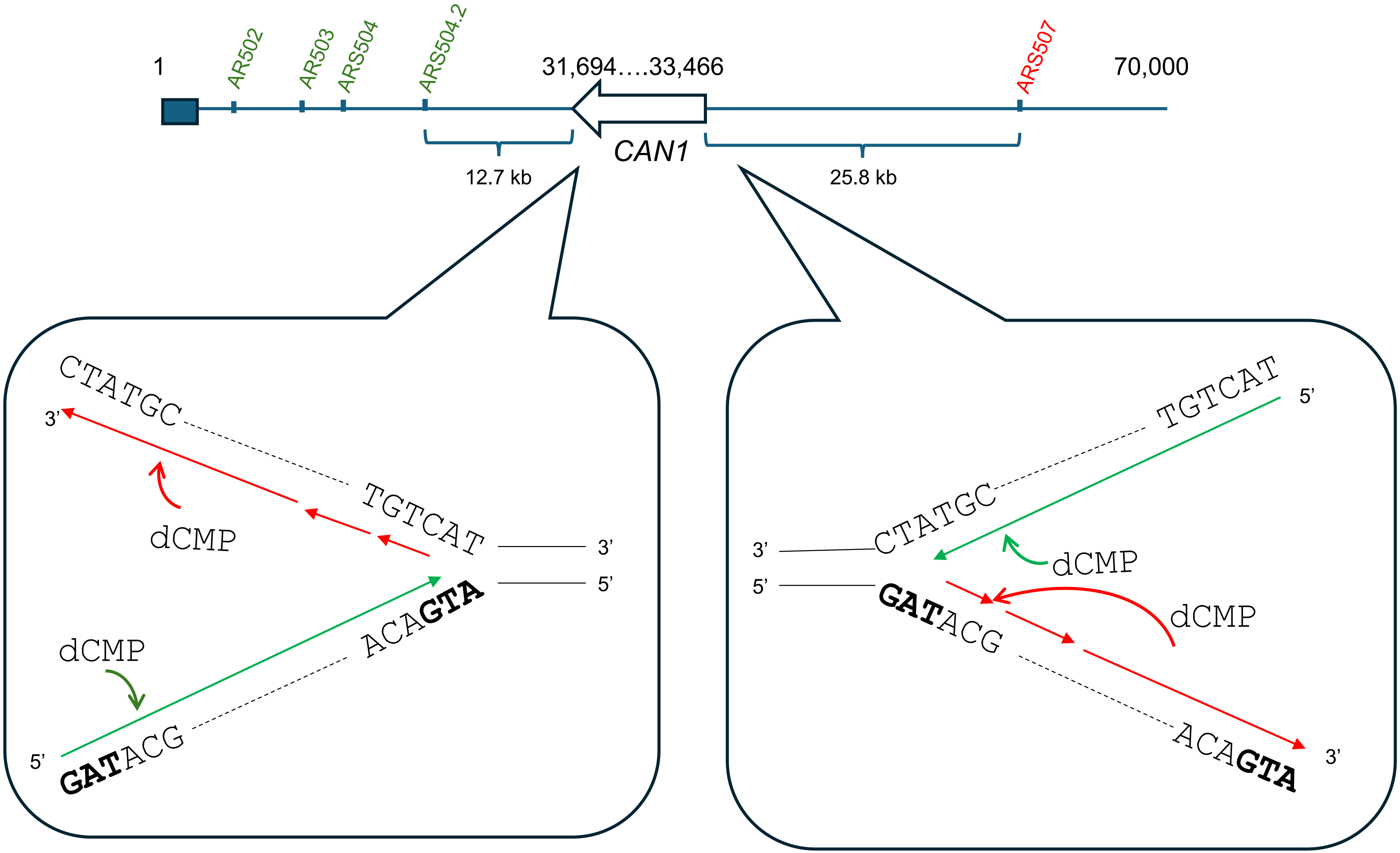

Supplemental Fig. 8. Schematic map of the left arm of chromosome V with positions of telomere (black box), ARS elements (small boxes) and the *CAN1* gene (open arrow showing start and end of the gene). Below is a replication forks passing *CAN1* and emanating from the left or to the right of the gene. dCMP misincorporations by Rev1 at the leading (green) and lagging (red) strands are shown. The diagram shows 6 nucleotides at start (initiating ATG in bold on non-transcribed strand) and end of the gene (terminating TGA in bold on non-transcribed strand). The sequence of the leading stans from right to left corresponds to the sequence  presented in sFig. 5.

**Supplemental File 1.**

*Identification of position-specific mutation hotspot candidates.*

Mutation hotspot candidates were identified assuming that mutations occur independently at each genomic position and follow a Poisson distribution under a homogeneous background mutation model for each nucleotide class GC (G or C) and AT (A or T) (Luria and Delbrück 1943; Lynch 2010). We estimated the expected mutation rate as the ratio of the corresponding nucleotide class occurrence to the total *CAN1* length. We calculated a one-sided p-value for each *CAN1* position using the survival function of the Poisson distribution (scipy.stats). To control for multiple testing, p-values were adjusted using the Benjamini–Hochberg false discovery rate (FDR) procedure (Benjamini and Hochberg 1995). We considered positions with FDR-adjusted p-values below 0.05 as candidate mutation hotspots.

*Identification of strain-specific mutation hotspots*

To further identify genomic positions contributing to differences in mutation spectra, pairwise strain comparisons were performed using the $G^2$ likelihood-ratio statistic (Piegorsch 2005).To localize candidate positions contributing to heterogeneity, we ranked all non-zero *CAN1* positions for the compared strains by their individual contributions to the total $G^2$ statistic, and recalculated the $G^2$ statistic after each removal until the remaining table satisfied the homogeneity criterion. We assessed the empirical critical value of the $G^2$ statistic under the homogeneity criterion using the 95$^{th}$ percentile of 10,000 Monte Carlo–permuted contingency tables. The overall stepwise decomposition strategy follows methods developed for identifying sources of heterogeneity in contingency tables and mutation spectra analyses (Piegorsch 2005). We then evaluated the significance of each candidate strain-specific hotspot using permutation testing. We used the absolute difference in mutation counts between strains as the test statistic. We generated the empirical null distribution of the test statistic using Monte Carlo permutation sampling that preserves marginal totals. This approach follows the logic described by Adams and Skopek and permutation-based inference frameworks (Adams and Skopek 1987; Good 2005). Multiple testing correction for site-level comparisons was performed using the FDR procedure

(Benjamini and Hochberg 1995). We considered positions with FDR-adjusted p-values below 0.05 as candidate mutation hotspots.

*Sequence context enrichment analysis*

To characterize event-weighted sequence context biases of single base substitutions (SBS), we constructed base aware enrichment logos centered on pyrimidines (C and T). For each strain, we extracted sequence contexts around mutated sites and reoriented them so all substitutions were represented as C- or T-centered events, analogous to the 96-channel mutational framework described by (Nik-Zainal et al. 2012). We visualized log-odds matrices as enrichment–depletion logos, following principles similar to the EDLogo framework proposed in (Dey et al. 2018). Sequence logo plots were generated using the Python package Logomaker (Tareen and Kinney 2020).

We calculated flanking nucleotide enrichment as position-specific log-odds ratios comparing nucleotide frequencies in hotspot-centered windows to *CAN1* background frequencies. For each flanking position, we median-centered log-odds values across nucleotides to reduce compositional bias and highlight relative enrichment patterns.

In contrast, we treated central nucleotides separately and reported raw (non-centered) log-odds values. We calculated central log-odds as the $\log_2$ ratio between the observed frequency of each central base (C or T) at hotspot positions and its corresponding gene frequency across all eligible positions in the analyzed sequence. This separation ensures that flanking positions reflect relative positional preferences, while central positions reflect overall enrichment or depletion of mutation categories.

*Single base substitution (SBS96) spectrum analysis*

We classified single base substitutions (SBS) into trinucleotide mutation contexts following the pyrimidine-centered convention (Nik-Zainal et al. 2012), yielding 96 possible mutation classes (6 substitution types × 16 trinucleotide contexts). For each strain, we aggregated mutation counts across the 96 SBS classes to construct mutation spectra. To account for sequence composition bias, we normalized trinucleotide frequencies by the corresponding trinucleotide frequency in *CAN1*. To evaluate similarity between experimentally derived normalized SBS96 vectors and previously reported COSMIC mutational signatures, cosine similarity was calculated (Alexandrov et al. 2013a). We assessed the statistical significance of similarity using Monte Carlo permutation. We calculated empirical p-values based on the proportion of permuted similarity values exceeding the observed similarity. We quantified pairwise similarity between strain-specific SBS96 spectra using cosine similarity. We estimated confidence intervals (CI) for cosine similarity using bootstrap resampling. To do this, we resampled mutations with replacement from the observed mutation counts for each strain to generate synthetic SBS96 spectra of equal total size. We normalized bootstrapped spectra using the same trinucleotide frequency correction applied to the original data. We computed cosine similarity for each pair of bootstrapped spectra across 1,000 bootstrap iterations, generating an empirical distribution from which we used the 2.5th and 97.5th percentiles to define the 95% confidence interval.

To identify underlying mutational components across strains, we decomposed normalized SBS96 spectra using non-negative matrix factorization (NMF), following approaches established for mutational signature analysis (Alexandrov et al. 2013b). The input matrix consisted of normalized SBS96 vectors from all analyzed strains, with each row representing a strain and each column corresponding to one of the 96 trinucleotide mutation contexts.

We set the number of components to two based on interpretability and the stability of the resulting spectra. We normalized the extracted components to sum to unity and compared them to reference mutational signatures from the COSMIC database using cosine similarity; we then visualized the resulting scores alongside the reference COSMIC signatures with the highest cosine similarity to facilitate qualitative comparison.
